# Noise source importance in linear stochastic models of biological systems that grow, shrink, wander, or persist

**DOI:** 10.1101/2022.01.10.475598

**Authors:** Alexander Strang, William Huffmyer, Hilary Rollins, Karen C. Abbott, Peter J. Thomas

## Abstract

While noise is an important factor in biology, biological processes often involve multiple noise sources, whose relative importance can be unclear. Here we develop tools that quantify the importance of noise sources in a network based on their contributions to variability in a quantity of interest. We generalize the edge importance measures proposed by Schmidt and Thomas [1] for first-order reaction networks whose steady-state variance is a linear combination of variance produced by each directed edge. We show that the same additive property extends to a general family of stochastic processes subject to a set of linearity assumptions, whether in discrete or continuous state or time. Our analysis applies to both expanding and contracting populations, as well as populations obeying a martingale (“wandering”) at long times. We show that the original Schmidt-Thomas edge importance measure is a special case of our more general measure, and is recovered when the model satisfies a conservation constraint (“persists”). In the growing and wandering cases we show that the choice of observables (measurements) used to monitor the process does not influence which noise sources are important at long times. In contrast, in the shrinking or persisting case, which noise sources are important depends on what is measured. We also generalize our measures to admit models with affine moment update equations, which admit additional limiting scenarios, and arise naturally after linearization. We illustrate our results using examples from cell biology and ecology: (i) a model for the dynamics of the inositol trisphospate receptor, (ii) a model for an endangered population of white-tailed eagles, and (iii) a model for wood frog dispersal.

**Author summary:** Biological processes are frequently subject to an ensemble of independent noise sources. Noise sources produce fluctuations that propagate through the system, driving fluctuations in quantities of interest such as population size or ion channel configuration. We introduce a measure that quantifies how much variability each noise source contributes to any given quantity of interest. Using these methods, we identify which binding events contribute significantly to fluctuations in the state of a molecular signalling channel, which life history events contribute the most variability to an eagle population before and after a successful conservation effort rescued the population from the brink of extinction, and which dispersal events, at what times, matter most to variability in the recolonization of a series of ponds by wood frogs after a drought.

## Introduction

*Noise source importance* evaluates the relative contributions of a set of noise sources to variability in a quantity of interest. By evaluating noise source importance we can quantify when, where, and to which quantities, individual noise sources matter.

Noise source importance was originally introduced to identify unimportant noise sources that could be ignored as a form of model reduction [1, 2]. In some cases, such as in the Hodgkin-Huxley model for neuron firing, this approach can lead to efficient, yet accurate, approximate simulation [3–5]. To illustrate this application we will ask, which binding events can be ignored when simulating the inositol triphosphate channel responsible for calcium induced calcium release?

More broadly, when fluctuations in a quantity are important, it is helpful to know where those fluctuations come from. For example, variance in the reproductive success of individual organisms can be decomposed into variability contributed by individual traits, and variability arising from random events during the life of the individual. The relative importance of these two sources determine whether an individual’s success is determined primarily by their intrinsic quality (pluck), or luck during their lifetime. Snyder et al. applied this decomposition to show the lifetime reproductive success of a shrub and perrenial grass (*Artemisia tripartita* and *Pseudoroegneria spicata*) are more determined by luck than the quality of their location [6].

The same kinds of questions can be asked for other noise sources and quantities of interest. In most conservation efforts, population size is the most important quantity. Variability in population size drives stochastic extinction for small populations, so controlling variability is important for their conservation. As an example, we will ask, did variation in clutch size, juvenile survival, or adult survival play a more important role in variability in a population of white-tailed eagles (*Haliaeetus albicilla*) while they were endangered? Have the important sources of variability changed now that the eagle populations have recovered? Fluctuations also play an important role in colonization and invasion processes triggered by rare dispersal events. We will ask, does variability due to dispersal play an important role in wood frog (*Rana sylvatica*) populations when at equilibrium, and during a recolonization process following a drought?

Note that the variables observed, or choice of measurements used to monitor a process, can strongly influence which noise sources are important. Indeed, noise source importance measures were originally introduced to answer the question: which noise sources are important *to what observables* [7]? This question is important in cellular signalling, where it is often only possible to measure whether a channel is open or closed, not its specific configuration. Similarly, in population biology it is often impossible to exhaustively survey all relevant variables. Then measurement choices must be made. These could be guided by identifying transitions that contribute significant variability to the quantity of interest. We would then ask: which noise sources are important *to what quantities of interest*? Thus, in addition to asking which noise sources are important, we will ask: how does the choice of observable influence which noise sources are important?

To address these questions, we extend an existing edge importance measure, introduced by Schmidt and Thomas for chemical reaction networks with a conservation constraint [1]. In chemical reaction networks, each edge in the reaction network is an independent noise source. Noise associated with a reaction arises from variability in the timing of reaction events, and can be removed from models by replacing the number of reaction occurrences in a time interval with the expected number of occurrences. Reducing the total number of noise sources in the model can allow more efficient simulation without changing the expected dynamics [2]. Removing noise sources usually reduces the variability in the process, so it is important to choose the set of edges simulated deterministically with care. If the reaction network consists entirely of first-order reactions, then the steady state variance in an affine measurable function of the state variables can be expressed as a sum of variance contributed by each edge in the network [1]. The affine measurable function models the observable or quantity of interest. The importance of each noise source to variability in the quantity of interest can be evaluated by computing the fraction of the total variability contributed by each edge. Often the important edges are edges which directly change the observable, while the observable is “shielded” from fluctuations on the other edges [2]. The stochastic shielding approximation reduces a reaction network by dropping noise from unimportant edges [1, 2, 7]. Example applications to neural dynamics are discussed in [3–5, 8–10].

The importance measure developed by Schmidt and Thomas is limited to stochastic models with a very specific structure. First order chemical reaction networks are discrete-state continuous-time Markov chains with linear propensities. Moreover, the signalling models considered by Schmidt and Thomas always satisfy a conservation constraint, since the state space models configurations of a fixed number of channels. As a result, the models necessarily approach a steady state distribution.

Many important biological models do not fit a reaction network framework, evolve continuously in time, use linear propensities, satisfy a conservation constraint, or possess a steady state distribution. For example, population models used to inform species management and conservation cannot assume a conserved population, because the principal modelling objective is to predict population growth or decline. Similarly, branching process models are widely used to study cell reproduction in oncology, and do not necessarily possess a steady state distribution as the population of cells may be expected to grow or shrink over time [11]. We will consider two representative population models (see §Applications) which do not fit the reaction network framework, do not occur continuously in time, and do not necessarily approach a steady state distribution, yet admit an edge-importance decomposition.

In this paper we introduce a general family of measures that provide such an edge-importance decomposition. Our generalization covers a broad class of stochastic models that satisfy a set of linearity assumptions. These include discrete- and continuous-state and time models, and models without a steady state distribution. The essential generalizations are listed below:

1. **Sufficient conditions**. We provide a set of sufficient conditions that a stochastic model must satisfy in order for the measures to be applied. By providing a set of minimal requirements, rather than an explicit model construction, we can address a much more general class of models, and establish results that hold independent of specific model formulations. We provide examples of common model formulations that satisfy these requirements, but future researchers could use the minimal requirements to check whether their models are amenable to our analysis.
2. **Generic asymptotic behavior**. Previous work was limited to models with a non-degenerate steady state distribution. We allow for models without a steady state, and whose expected state could diverge, approach a non-zero value, or converge to zero. We show that the existing measure is a special case of our general measures, and is recovered when the model satisfies a conservation constraint.
3. **Importance at finite times**. Many models are not globally linear, but can be approximated locally with a linearized model. Long-time limits of linear approximations may not be meaningful if the process is expected to leave the region where the linear approximation is valid. Then it is useful to compute noise source importance at intermediate times. Moreover, in some cases we are interested in the variability along a transient trajectory. For example, when considering an invasion process, we are typically interested in dynamics during the transient approach to equilibrium, not at equilibrium.

The paper is organized as follows. In §Scope we outline the sufficient conditions required for a model to suite the analysis. These conditions define the scope of models considered. We then present some important model classes that satisfy the sufficient conditions.

We divide our subsequent results into theory and application. In §Theoretical Results we derive the noise source importance measures for discrete- and continuous-time models, at finite and long times, in each possible limiting scenario, and show that Schmidt and Thomas’ measure [1] is a special case of our generalized measure. We demonstrate that the measures all share a generic derivation in terms of the production and propagation of variance, and the distinctions between the measures arise from whether most present variability was produced recently, or in the far past. We then compute the long time limit of the important measures in each limiting scenario, and find that, in contrast to the existing literature, when the process grows or wanders, long time noise source importance is independent of the choice of observable (c.f. Lemma). We conclude our theoretical investigation by presenting the equivalent measures in the affine case.

In §Applications we apply our theoretical results to illustrative examples corresponding to sample model types introduced in §Scope. The applications illustrate how the noise importance measures can be computed for widely used classes of stochastic models. We consider applications to cellular signalling to demonstrate consistency with existing methods in a model system where noise source importance has not been computed before. A reader interested in efficient simulation and model reduction can skip to §Inositol Trisphosphate and Calcium Signalling on page 15. We also consider applications to white-tailed eagles to demonstrate that our methods can be applied to discrete-time matrix models without a steady state population, and to a nonlinear model of wood frog dispersal to illustrate how our methods can be applied to linear approximations both about equilibrium and along transients associated with recolonization. A reader interested in conservation and variability in natural populations can skip to §White-Tailed Eagles (*Haliaeetus albicilla*) on page 21 or §Wood Frogs (*Rana sylvatica*) on page 24.

## Scope

### Sufficient Conditions

Let *X*(*t*) denote a continuous-time stochastic process and *X*_*t*_ a discrete-time stochastic process. We require the following properties:

P1 **Markov:** The stochastic process is Markovian.

P2 **Linearity of Conditional Expectation:** If *X*(*t*) is a continuous-time model, then 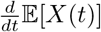, given *X*(*t*) = *x*, is either an affine or linear function of *x*. If *X*_*t*_ is a discrete-time process, then 𝔼[*X*_*t*+Δ*t*_], given *X*_*t*_ = *x*, is an affine or linear function of *x*.

P3 **Linearity of Conditional Variance:** If *X*(*t*) is a continuous-time process, then 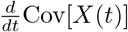, given *X*(*t*) = *x*, is either an affine or linear function of *x*. If *X*_*t*_ is a discrete-time process, then Cov[*X*_*t*+Δ*t*_], given *X*_*t*_ = *x*, is an affine or linear function of *x*.

P4 **Noise Separability:** If *X*(*t*) is a continuous-time process, then 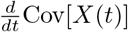 can be expressed as a sum over noise sources, the derivative is well-defined (symmetric and positive-definite) for any partial sum over the noise sources, and the derivative is zero if no noise sources are included. If *X*_*t*_ is a discrete-time process then Cov[*X*_*t*+Δ*t*_] can be expressed as a sum over noise sources, the covariance is well defined (symmetric and positive-definite) for any partial sum over noise sources, and the covariance is zero if no noise sources are included. In either case the contribution from each noise source is an affine function of *x*.

The first three conditions ensure that the state covariance at time *t* takes a canonical form. The last condition ensures that the state covariance can be separated into contributions from each noise source. The results provided in the main text assume that both the conditional expectation and variance are strictly linear functions of the state *x*. The generalization to affine functions of *x* is provided in S3 Appendix.

### Example Model Types

The following models satisfy the sufficient conditions:

#### Continuous-Time First-Order Reaction Networks

A continuous-time first order reaction network is specified by a set of reactions ℛ, and a propensity *λ*_*r*_(*x, t*) and stoichiometry *s*_*r*_ for each reaction *r* ∈ ℛ. The propensity *λ*_*r*_(*x, t*) is the expected rate at which reaction *r* occurs at time *t*. In a first order reaction network, the propensities are linear functions of *x*, denoted 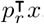. The actual reaction times are random. The probability that reaction *r* occurs once in a small time interval is *λ*_*r*_(*x, t*)Δ*t* + 𝒪 (Δ*t*^2^), doesn’t occur is 1 − *λ*_*r*_(*x, t*)Δ*t* + 𝒪 (Δ*t*^2^), and occurs more than once is 𝒪 (Δ*t*^2^). If reaction *r* occurs at time *t* then *X*(*t*) is replaced with *X*(*t*) + *s*_*r*_.

#### Langevin Approximation to Continuous-Time First-Order Reaction Networks

The Langevin approximation to a continuous-time first order reaction network is *dX*(*t*) = *AX*(*t*)*dt* + *B*(*x, t*)*dW*(*t*) where *W*(*t*) is a Wiener process, 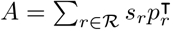 and 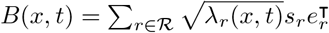 where *e*_*r*_ is the *r*^*th*^ |ℛ| × 1 indicator vector [7, 12, 13]. The associated diffusion matrix, 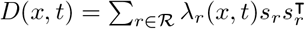, governs the instantaneous production of variance, while *A* governs the expected behavior of *X*(*t*).

#### Discrete-Time First-Order Reaction Networks

A discrete-time first order reaction network is also specified by a set of reactions, propensities which are linear functions of *x*, and stoichiometries. However, the number of times each reaction occurs in each time interval is Poisson distributed with mean *λ*_*r*_(*X*_*t*_, *t*)Δ*t* (c.f. [2]).

#### Discrete-Time Matrix Models

A discrete-time matrix model is a model where the distribution of *X*_*t*+Δ*t*_, given *X*_*t*_ = *x*, is parameterized by a linear function of *x*, say *Ax* for some matrix *A*, and some fixed external parameters. Usually the expected value of *X*_*t*+*δt*_ is given by *Ax*. For example, we could set each entry of *X*_*t*_ equal to a Poisson random variable, with means fixed by *AX*_*t*−Δ*t*_.

In practice, not all discrete-time matrix models use Poisson distributed random variables. Instead, the product *a*_*ij*_*x*_*tj*_ may represent the mean of a different distribution. In an age or stage-structured population model, the entries of the first row of *A* represent fecundity rates (reproduction), and the remaining entries all represent transition probabilities between stage/age classes *j* and *i* [14]. Then, the actual number of individuals who transition from class *j* to class *i* is binomially distributed, with *x*_*tj*_ trials and probability *a*_*ij*_. Thus the expected number of individuals making the transition is *a*_*ij*_*x*_*tj*_, but the variance in the number is *a*_*ij*_(1 −*a*_*ij*_)*x*_*tj*_, not *a*_*ij*_*x*_*tj*_. Nevertheless, the variance remains a linear function of *x*. Transitions representing survival, dispersal, aging, or growth often take this form. Binomial survival distributions appear widely in discrete time population models subject to demographic stochasticity [15].

Fecundity distributions are more model dependent, and may not be known explicitly [16]. While some authors and software packages adopt the Poisson distribution [17, 18], the Poisson distribution is often ill-suited to modeling the distribution of offspring since it often lacks biological justification, is unbounded above, and may have unrealistic variance given its mean. While a different parametric family of distributions could be used, for example, the generalized Poisson recommended in [16], our analysis depends only on the functional form of the conditional moments (see §White-Tailed Eagles). If it is assumed that, conditioned on their shared environment, the number of offspring produced by any individual (or mating pair) is independent of the number produced by any other individual, then the variance in the number of offspring produced by any age or stage class is a linear function of the number of individuals in that class. Thus it is enough to know the variance in the number of offspring per individual or mating pair to perform our analysis.

## Results

### Theoretical Results

If the stochastic process satisfies conditions P1-P4 above, then the state covariance takes a standard form. Let *V*(*t*) or *V*_*t*_ denote the covariance in the state, and 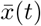 or 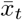 denote the expected state, at time *t*. Here we provide an explicit derivation for *V*_*t*_ given 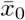 and *V*_0_. For simplicity, we assume that the time units are chosen so that Δ*t* = 1. The results for continuous-time systems are entirely analogous. We articulate the explicit forms for each case. Appendix S1 Appendix provides the derivation for continuous-time systems. In this section we restrict attention to the case when the conditional expectation and variance are strictly linear functions of *x*. The affine case is treated in S3 Appendix, and results are reported at the end of that section.

Under conditions P1-P2, the expected state given the previous state is a time-independent linear or affine function of the previous state. In the linear case there exists some update matrix *A* such that:

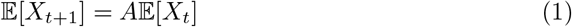

in which case, 𝔼[*X*_*t*_] = *A*^*t*^𝔼[*X*_0_].

The covariance also satisfies a recursion. Since the process is Markovian, we can apply the law of total variance to decompose the covariance at time *t* + 1:

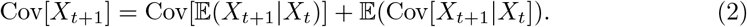

The expected value of *X*_*t*+1_ given *X*_*t*_ is *AX*_*t*+1_ so:

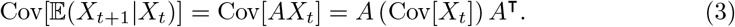

This term represents the forward propagation of past uncertainty into the present. The product *A*(·)*A*^⊤^ is the forward propagation operator.

Under condition P3, there exists some linear function of *x, D*(*x, t*), such that:

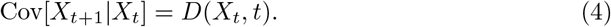

Then, since *D*(*x, t*) is linear:

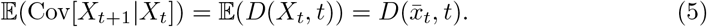

Therefore the covariance satisfies the recursive update equation:

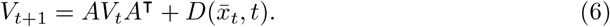

The recursion closes, leaving:

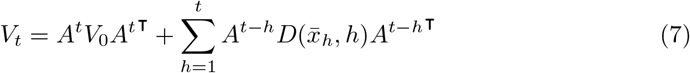

where 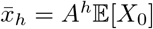. The proof follows by induction. The first term in Eq. 7 is the forward propagation of uncertainty in the initial conditions to the variance at time *t*, while the second term is the forward propagation of the additional uncertainty introduced at each intermediate time step. Each term in the sum is the uncertainty produced at time 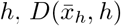, carried forward by the propagation operator 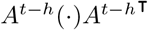.

The state covariance for a continuous-time process takes an analogous form:

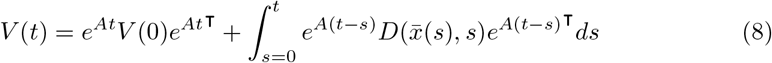

where *A* is the matrix such that 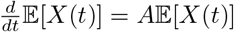 and *D*(*x*(*t*), *t*) is the matrix such that 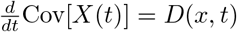 if *X*(*t*) = *x*. In continuous time the forward propagation of uncertainty is governed by 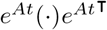, and uncertainty is produced continuously, so the sum over the past is replaced with an integral.

In both cases, the second term is the variance produced by the process during the time interval [0, *t*]. We will generally focus on this term, since the variance inherited from uncertainty in the initial conditions is independent of the noise sources. The variance produced by the process with initial expectation 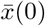 is the same as the variance in the process if initialized at 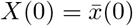. Therefore, for the rest of this discussion we drop the initial variance from Eq. 7 and Eq. 8.

#### Noise Source Expansion and Importance

Under condition P4, we can decompose the variance production term *D*(*x, t*) into a sum over noise sources. Let 𝒩 denote the set noise sources, and let *D*_*n*_(*x, t*) be the variability in *X*_*t*_ or *X*(*t*) produced by the *n*^*th*^ noise source. Then:

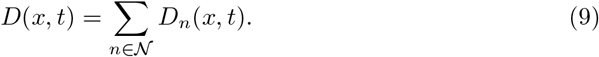

The variance produced by the process is:

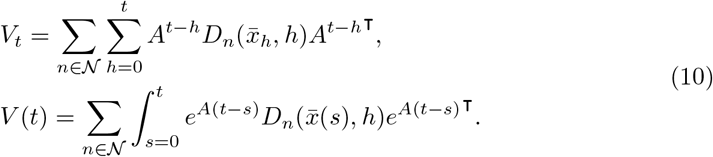

Two important properties of the state covariance can be inferred directly from Eq. 10. First, the uncertainty at any time *t* can be expressed as a sum over each noise source at each time in the past. In a simulated process, this decomposition expresses the uncertainty at time *t* as a sum over uncertainty contributed by each previous random number draw. It follows that the state covariance is additive in the sense that:

##### Lemma 2: (Additivity)

*If* 𝒮(*t*) *and* 𝒮′(*t*) *are disjoint subsets of* 𝒩 *at all t, then the state covariance in the process with noise sources* 𝒮 ∪ 𝒮′ *equals the sum of the covariances in the reduced processes with noise sources* 𝒮 *and* 𝒮 ′ *separately*.

The second property of eq. 10 involves ordering of covariance matrices. Covariance matrices can be (partially) ordered since they are real-symmetric and positive semi-definite. If *B* and *C* are both Hermitian positive-definite then *B* ≥*C* if *B* − *C* is positive semi-definite, and *B* > *C* if *B* − *C* is positive definite [19]. If || · ||is a monotone matrix norm then *B* ≥ *C* implies ||*B*|| ≥ ||*C*||. From here on || · ||will denote a monotone matrix norm.

Eq. 10 ensures that the state covariance is monotonically nondecreasing as more noise sources are added. The noise source expansion in Eq. 10 represents the variance produced by the process as a sum of positive semi-definite matrices, one for each noise source. Therefore:

##### Lemma 2: (Monotonicity)

*If* 𝒮′(*t*) ⊆ 𝒮(*t*) *for all t and V*′(*t*) *and V*(*t*) *are the covariances produced by the processes with noise sources* 𝒯(*t*) *and* 𝒮′(*t*) *then V*′(*t*) ≤ *V*(*t*) *and* ||*V*′(*t*)|| ≤ ||*V*(*t*)|| *for any monotone matrix norm*.

The importance of a set of noise sources can be analyzed by considering the variance produced by a reduced process with the noise removed from the remaining sources. Let 𝒮(*t*) denote the set of noise sources that are included in the process at time *t*. Then the variance produced by the reduced process is given by restricting the sum over noise sources to the sum over 𝒮.

To analyze noise source importance, we also need to specify how we measure the size of a covariance matrix, and whether we are interested in the full state of the process or some observable function of the state. Here we restrict our focus to observables that are linear functions of the state. Let *M* ^⊤^ be the Jacobian of the observable so that each column of *M* corresponds to a specific observable, and inner products with columns of *M* represent measurements. The matrix *M* is the measurement matrix. Then the importance *R* of noise sources 𝒮 at time *t*, given initial expectation 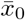, and measurement matrix *M*, is the ratio of the norm of the covariance in the observable for the reduced process, to the norm of the covariance in the observable for the full process. That is, if 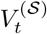 denotes the state covariance in the process with sources 𝒮, then:

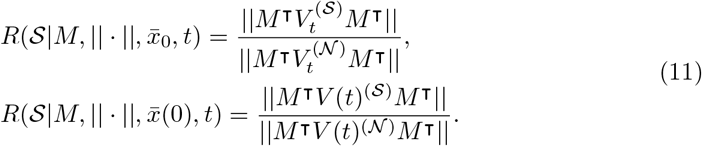

Then, for discrete and continuous-time respectively:

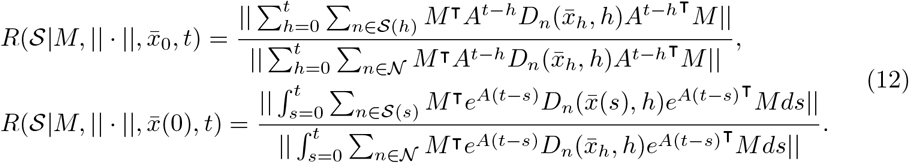

Note that if the chosen norm is linear in the matrix entries, then the importance can be interpreted as the fraction of the uncertainty contributed by noise sources 𝒮. It is also natural to seek a matrix norm that is unitarily invariant, so that the size of the covariance is independent of our representation of the state space. The only linear, unitarily invariant matrix norm on symmetric-positive definite matrices is the trace norm. The trace is a monotone norm for positive semi-definite matrices since the diagonal entries of a positive semi-definite matrix are all nonnegative. The trace is also a natural choice in this context, since the trace of a covariance is the expected (squared) distance between a sampled observable and its expectation.

#### Eigen-Expansion and Closed Form

Suppose that the matrix *A* is diagonalizable and the set of noise sources considered, 𝒮(*t*), is constant in time. Then we can close the sum/integral over the past. The discrete-time derivation is provided here.

If *A* is diagonalizable then *A* = *U* Λ*W* ^⊤^ where *U* and *W* are matrices whose columns are the right and left eigenvectors respectively, and *W*^⊤^ = *U* ^−1^. Then:

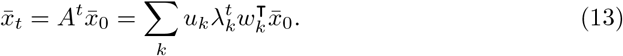

Next, since *D*_*n*_(*x, t*) is linear in *x*, and time independent:

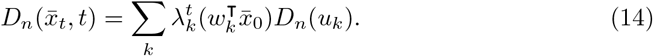

Therefore, the variance produced by the process at time *t* is:

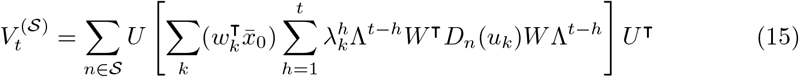

where the sum over noise sources can be moved outside the sum over time via the assumption that the set of noise sources is unchanging.

Next, define the matrix:

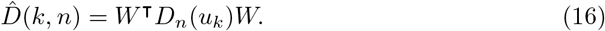

Then Eq. 15 can be expanded into a sum over triples of eigenvalues:

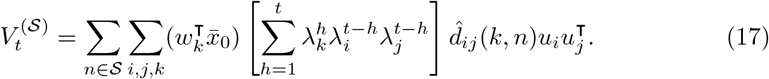

The sum can be rewritten as a geometric series, so:

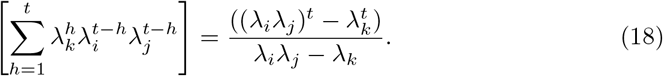

In the special case when *λ*_*i*_*λ*_*j*_ = *λ*_*k*_ the sum equals *λ*_*k*_^*t*^*t*.

Therefore, the closed form for the covariance produced by the reduced process with noise sources 𝒮 is:

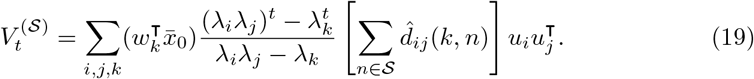

The analogous equation in continuous-time is:

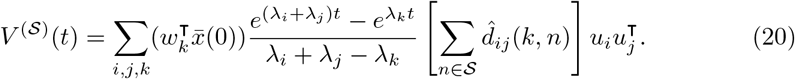

Then, to compute finite time importance, substitute Eq 19 or 20 into Eq 11. Note that the existing measure introduced by Schmidt and Thomas, only considered importance in the long time limit [1]. Finite time importance is of interest in applications whenever fluctuations about transients matter, such as during an invasion process.

The only difference in the discrete and continuous-time expressions is the time independent term in *V* ^(𝒮)^(*t*) involving the eigenvalues in equations 19 and 20. If the discrete-time model is chosen to discretize a continuous-time model then the two expressions converge in the limit of small Δ*t* (see S2 Appendix).

#### Long-Time Limits

Eq. 11 establishes a closed form for finite-time noise source importance. The long-time limit remains.

For processes with a non-degenerate steady state distribution, the state covariance converges to a fixed matrix. For processes without a steady state, the covariance can diverge or converge to zero. In these cases, we scale the covariance by its asymptotic growth rate, and analyze importance with respect to the scaled covariance. Scaling the variances by the asymptotic growth rate does not change the importance measure since ||*αB*|| = |*α*| · ||*B*||, so the numerator and denominator are scaled by the same constant.

To analyze the long-time limit we require an additional property:

P5 **Dominant Eigenvalue:** *A* has a dominant eigenvalue *λ*_1_ such that |*λ*_1_| > |*λ*_*j*_|, in discrete time, or Real(*λ*_1_) > Real(*λ*_*j*_), in continuous time, for any *j* > 1.

The limiting behavior of the variance depends on the limiting behavior of Eq. 18 as *t* gets large. These limits depend, in turn, on whether |*λ*_1_| is greater than, equal to, or less than one, which determines whether *X*_*t*_ is expected to grow, remain constant, or shrink. When growing, the variance grows proportionally with 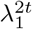, when neutral the variance grows proportionally with *t*, and when shrinking the variance shrinks proportionally with 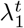.

The respective limits are:

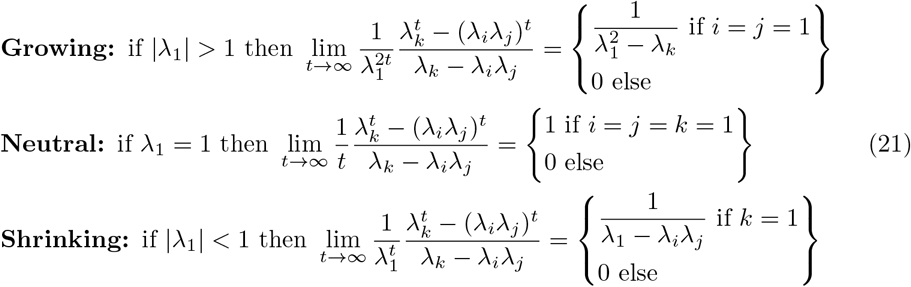

In each case, the scaled variances take the general form:

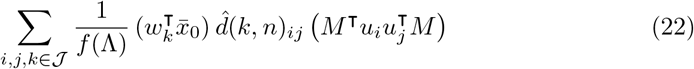

where 𝒥 is an index set and *f* (Λ) is a function of the eigenvalues. Both 𝒥 and *f* (Λ) depend on the limiting scenario, cf. Table 1.

**Table 1.**
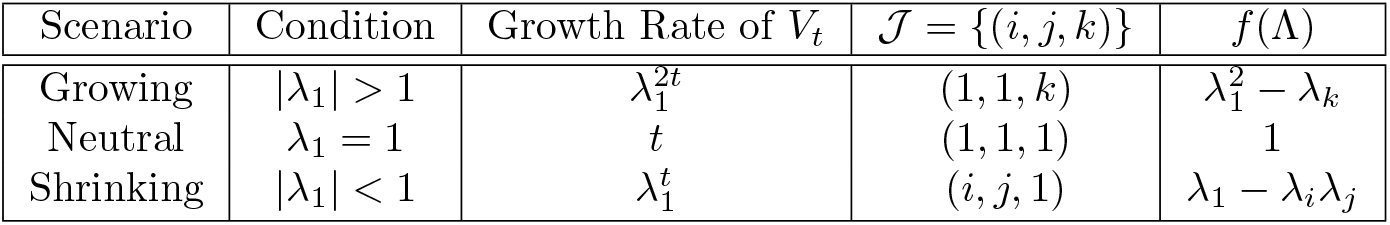
Index set and eigenvalue dependence for long time limit of discrete-time variances, scaled by their growth rate.

Note that each of the cases has different limiting behavior. For example, when growing, the covariance converges to a constant when rescaled by 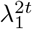, the growth rate of 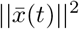. Then the limiting behavior of the scaled covariance is equivalent to the covariance in 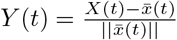, the relative fluctuations about the expected state. When shrinking, the variance is proportional to 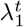 instead of 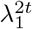. The difference in these limiting behaviors can be explained by considering the primary sources of uncertainty at long times.

If *X*(*t*) is expected to grow, then the expected trajectories diverge exponentially. As a result, the uncertainty at some long time is mostly generated by uncertainty produced at the start of the process. Thus, uncertainty early in the process (far in the past) is compounded over time, amplified by the diverging trajectories.

In contrast, if *X*(*t*) is expected to shrink, then all the expected trajectories are converging exponentially. As a result, the uncertainty at some long time is mostly generated by fluctuations in the immediate past. The influence of uncertainty early in the process (far in the past) is suppressed as the trajectories converge.

To see this distinction mathematically, consider the contribution of variance introduced at time *h* to the state variance at time *t* > *h*. This contribution depends on 2*t* −*h* products with the matrix *A*, since the expected state at time *h* depends on *A*^*h*^, and the forward propagation of variance depends on a product on the left and right with *A*^*t*−*h*^. If some of the eigenvalues of *A* are greater than one, then maximizing the exponent 2*t* −*h* will maximize products with *A*^2*t*−*h*^. For fixed *t*, the exponent is maximized for *h* = 0 (the distant past) and the variance grows in proportion to 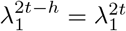. In contrast, if all of the eigenvalues of *A* are less than one, then minimizing the exponent 2*t* −*h* will usually maximize products with *A*^2*t*−*h*^. The exponent is minimized for *h* = *t* (the near-term) and the variance decays in proportion to 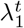. Thus, the distinction between the growing and shrinking cases comes from whether the dominant contribution to the long time variance is the amplification of past uncertainty, or uncertainty generated in the immediate past.

What if *X*(*t*) is neither expected to grow nor shrink? Then there is a direction along which *X*(*t*) is a martingale. Specifically, if *λ*_1_ = 1, then the component of *X*(*t*) along the dominant eigenvector, *u*_1_, is an unbiased random walk. This random walk accrues variance linearly in time. Past variance does not decay or grow, so the variance along *u*_1_ grows steadily with each added step. Meanwhile, the projection of *X*(*t*) onto all other eigenvectors vanishes, so the long time variance is dominated by the linear growth along *u*_1_.

The continuous-time results are entirely analogous. In fact, Eq. 22 applies, with the same choice of index set, only with different *f* (Λ). The necessary choices are summarized in Table 2.

**Table 2.** Index set and eigenvalue dependence for long time limit of continuous-time variances, scaled by their growth rate.

| Scenario | Condition | Growth Rate of $V(t)$ | $\mathcal{J} = \{(i, j, k)\}$ | $f(\Lambda)$ |
| --- | --- | --- | --- | --- |
| Growing | $\text{Real}(\lambda_1) > 0$ | $e^{2\lambda_1 t}$ | $(1, 1, k)$ | $2\lambda_1 - \lambda_k$ |
| Neutral | $\lambda_1 = 0$ | $t$ | $(1, 1, 1)$ | 1 |
| Shrinking | $\text{Real}(\lambda_1) < 0$ | $e^{\lambda_1 t}$ | $(i, j, 1)$ | $\lambda_1 - \lambda_i - \lambda_j$ |

The finite time noise source importance depends on the set of noise sources considered, the measurement matrix (the choice of observable), the chosen norm, and the initial expected state of the process. Our next Lemma answers the question: *What does the long time noise source depend on?*

##### Lemma 3

*In the growing case, the long time noise source importance*, 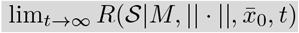, *is independent of the choice of measurement matrix M and norm*|| · ||. *In the shrinking case, the long time noise source importance is independent of the initial conditions* 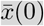. *In the neutral case without a conservation constraint, the long time noise source importance is independent of the choice of measurement matrix M, the norm* || · ||, *and the initial conditions*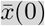.

**Proof:** In both the growing and neutral cases (without conservation constraint) all contributions to the covariance take the form of a scalar coefficient times 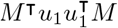. It follows that the variance associated with a set of noise sources is the sum of those scalar coefficients times the matrix 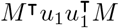. The same holds when using all noise sources. Therefore the covariance in any reduction of the process is proportional to the covariance in the full process. The ratio of the norm of two proportional matrices is the ratio of their proportionality constants, since ||*αA*|| = |*α*|||*A*|| for any norm. Thus, the asymptotic importance in the growing and neutral case only depend on the scalar coefficients that scale 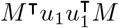.

In particular, in the growing case, the asymptotic importance for discrete and continuous models takes the form:

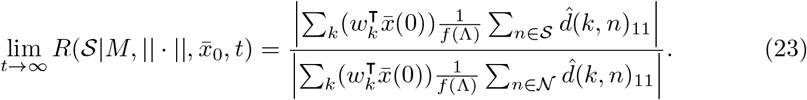

In the shrinking and (unconstrained) neutral cases, all contributions to the covariance are proportional to 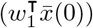, so the dependence on initial conditions cancels.

Thus, in the shrinking case:

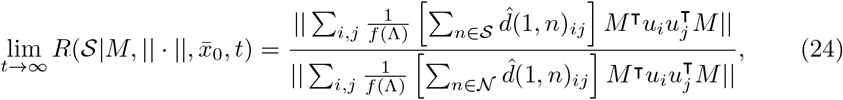

and in the unconstrained neutral case:

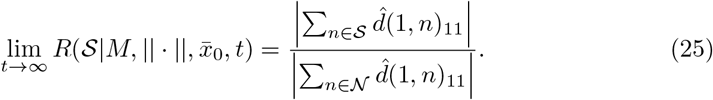

Therefore in the growing case the importance is independent of the measurement matrix and choice of norm, in the shrinking case the importance is independent of the initial conditions, and in the unconstrained neutral case the importance is independent of the measurement matrix, norm, and initial conditions. ■

Lemma establishes that noise source importance is independent of what is observed when the process grows or wanders. This result stands in contrast to the existing literature, where the choice of measurement strongly influences which noise sources are important. Indeed, the stochastic shielding heuristic as originally proposed by Schmandt and Galan [2] is predicated on the assumption that the only important noise sources are those that directly change the observable. In that case, edge importance is entirely determined by what is measured. Schmidt and Thomas introduced their measure to show that the heuristic does not always hold [1]. Nevertheless, edge importance reversal usually occurs when the dynamics of the observable are strongly coupled, albeit indirectly, to a distant noise source [7]. Thus, importance is still directed by the observable.

As Lemma **??** establishes, the long time noise source importance does not depend on the choice of observable, when the triple sum over *i, j, k* is restricted to a sum over *k*. Recall that the sum over *k* accounts for the expected dynamics up to an intermediate time, while the sum over *i, j* account for the propagation of variance after that intermediate time (cf. Eq. (17)). This occurs when most of the variance is produced in the distant past, as in the growing case, or when it accrues linearly along one direction and shrinks along all others, as in the wandering case. Note that, in both cases, the *finite* time importance still depends on the choice of measurement matrix. If the eigenvalues of *A* are ordered in decreasing magnitude, then the dependence on the measurement matrix decays as 𝒪 ((||*λ*_2_||*/*||*λ*_1_||)^*t*^) in the growing case, and as 𝒪 ((||*λ*_2_||^*t*^)*/t*) in the wandering case.

#### The Steady State Case

So far we have not considered a limiting scenario in which the process approaches a steady state distribution other than a delta distribution. The edge importance measure introduced in [1] is only defined for processes with a non-degenerate steady state distribution. The non-degenerate steady state case can be recovered from our analysis by enforcing a conservation constraint. This special case is recovered by the additional assumptions:

1. **Neutral:** |*λ*_1_| = 1 (discrete-time) or *λ*_1_ = 0 (continuous-time)
2. **Conservation:** The dominant left eigenvector *w*_1_ is in the nullspace of *D*_*n*_(*x, t*) for all *n, x, t*.

This situation occurs naturally in reaction networks where all the reactions have stoichiometry vectors which conserve a quantity (for example, the total number of particles or channels). The conservation constraint ensures that no noise source produces variance along *w*_1_, so the inner product of 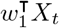 is conserved.

Then 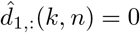 and 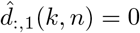, since *w*_1_ is orthogonal to the range of *D*, so any term in the triple sum in Eq. 17 with *i* = 1 or *j* = 1 is automatically zero, which sets the diverging term to zero. If *k* ≠ 1 then the entire term converges to zero, so the only surviving terms are *k* = 1, *i* ≠ 1, *j* ≠ 1. Then the long time limit (steady-state) covariance equals:

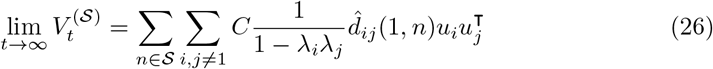

where 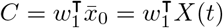 is the conserved quantity.

The analogous result in continuous-time is:

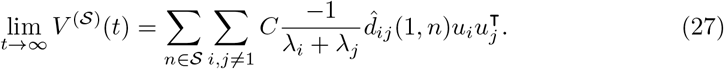

To compare to the edge importance measure proposed in [1] we write out 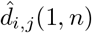 in detail. First, note that 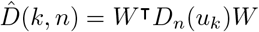. Next, if the continuous-time model is a reaction network, then each reaction *r* is a noise source, so we replace *n* with *r* and 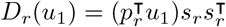. Then:

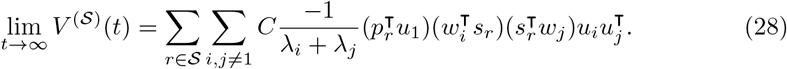

In a continuous-time first-order reaction network *u*_1_ is proportional to the steady-state distribution, and 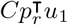 is the rate at which reaction *r* occurs at steady state. This flux, *J*_*r*_, is the steady state flux of reaction *r*. Then the uncertainty contributed by a single reaction is:

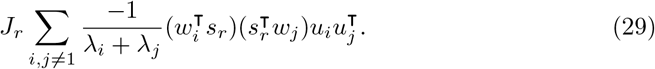

The associated asymptotic importance is:

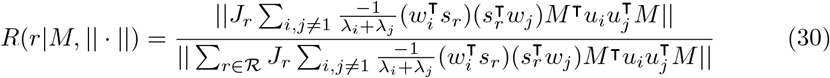

which coincides with the edge-importance measure defined by Schmidt and Thomas for first-order continuous-time reaction networks when the observable is one-dimensional (*M* has only one column) [1].

### Affine Conditional Moments

The previous sections assumed that the conditional moment equations were linear functions of the state. Here we generalize our results to the affine case. The derivation largely follows from the linear case and is provided in S3 Appendix.

In the affine case Eq. 1 and Eq. 4 are replaced with:

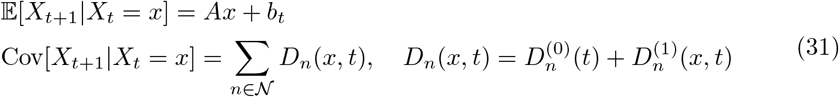

where each *D*_*n*_ (*x, t*) is an affine function of the state 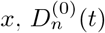 is the *x* independent part of each noise source (zeroth order term), and 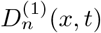 is the *x* dependent part of each noise source (first order term). Note that linearity requires both the conditional expectation and variance to be linear functions of *x*, which requires *b* = 0 and *D*^(*n*)^ = 0.

Affine conditional moments arise naturally in problems with source reactions. For example, consider a population subject to immigration from a reservoir not included in the state. Then the immigration process is independent of the state of the system, so adds state-independent terms to the conditional moments. Similarly, in a reaction network with source reactions, the source reactions randomly add new particles from a state-independent distribution, so the resulting conditional moments include state-independent terms.

Affine conditional moments also arise naturally when linearizing nonlinear stochastic models. If realizations of a nonlinear stochastic model tend to remain near some non-zero equilibrium state, then, provided the fluctuations are small relative to the second derivative of the reaction rate functions, we can approximate the stochastic dynamics by linearizing the rate functions, or by linearizing the otherwise nonlinear conditional moments. Linearization will generally produce affine conditional moments since, if the linearization is based on Taylor expansion, the zeroth order terms in the Taylor expansion will become state-independent terms in the conditional moments. Alternatively, if the linearization is based on a best fit linear model to observed dynamics then the linearization will include state-independent terms whenever the fit intercepts are nonzero. We consider an example of this kind in §Wood Frogs.

If the conditional moment equations are affine, then the moments satisfy the same recursions as before,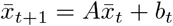 and 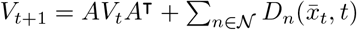. The recursions are unchanged since they depend only on the linearity of expectation, law of total variance, and linearity of the conditional moments.

While the recursions are unchanged, the long time limits of the moments are changed by adding state independent terms. In particular, adding constant source terms to the conditional moment equations introduces a new limiting scenario.

In absence of a constant source term, the only limiting scenario with a non-degenerate steady state distribution was the neutral case, *λ*_1_ = 1, with a conservation constraint. In the shrinking case, |*λ*_1_| < 1 the steady state distribution was a delta distribution at the origin. In the presence of constant source terms the |*λ*_1_| < 1 case admits non-degenerate steady states. If *b* > 0 and |*λ*_1_| < 1, then the expected state will converge towards a nonzero equilibrium, and thus the variance source terms will approach constant, nonzero values. The resulting steady state distribution balances the fluctuations produced by the nonzero noise sources, with their tendency to decay back towards the equilibrium. This is the type of steady state commonly observed in Ornstein-Uhlenbeck processes. If *b* = 0 then 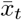 will converge to the origin. In the linear case the intensity of each noise source is proportional to *x*, so when 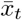 approaches the origin all the noise sources stop producing noise - hence the approach to a delta distribution. In the affine case, when *b* = 0 it must be true that 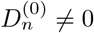, so the noise sources *D*_*n*_(*x, t*) do not vanish. Then the constant terms in the noise sources produce fluctuations even when the expected state approaches the origin. These fluctuations maintain a non-degenerate steady state.

As in the linear case, we can compute the long-time variance in the state by expanding the recursions onto the eigenvectors of *A*, then closing the resulting sums over the past, and evaluating the limiting behavior. Results for each limiting scenario are provided below (see S3 Appendix for derivation):

**Growing:** if |*λ*_1_| > 1 then:

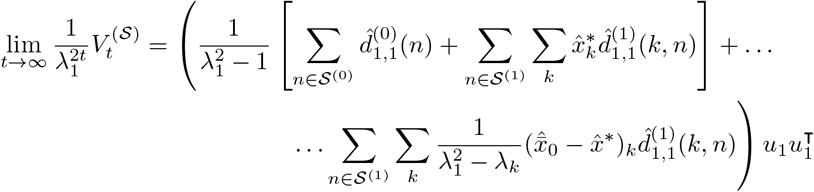

**Wandering:** if *λ*_1_ = 1 then:

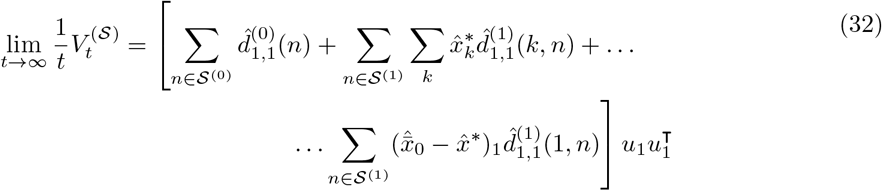

**Steady State:** if |*λ*_1_| < 1 then:

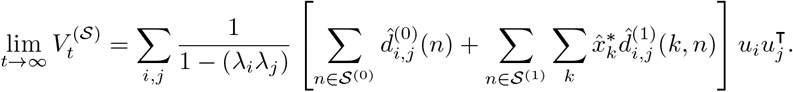

Here 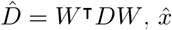 denotes expansion on the eigenbasis, 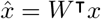, and *x*\* denotes the equilibrium of the expected dynamics, 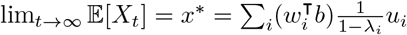.

Once the long time variance has been computed, the importance of each noise source can be analyzed using the usual approach. Computing the long time variance in the affine case is considerably more expensive than in the linear case, as all scenarios require computing a sum over each eigenvalue, and the steady state scenario requires computing a sum over all triples of eigenvalues.

### Sample Applications

The four models introduced in §Scope differ only in the formulation of *A* and *D*_*n*_(*x*). For both continuous-time reaction networks 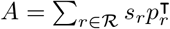 and 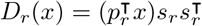. For a discrete-time reaction network with time step *τ* the matrices are 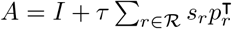 and 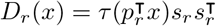. For a discrete-time matrix model, *A* is explicitly fixed by the model parameterization, while *D*_*n*_(*x, t*) will depend on the distributions used to sample the updates.

We consider three applied examples in detail. The first is a chemical reaction network model of a component of a cellular signalling process. We use this example to demonstrate the utility of the existing edge importance measure. The remaining two examples are structured population models, that demonstrate the extension to discrete-time, discrete-state models, with or without a steady state, evaluated at finite times. The last case demonstrates the importance of the affine generalization. Together, these studies show how to answer the question: *Which sources of noise are important to which quantities of interest?*

### Continuous-time Reaction Networks: Inositol Trisphosphate and Calcium Signalling

Cells utilize a variety of cellular signalling pathways to respond to external stimuli and to perform essential functions. Calcium ions, Ca^2+^, are widely used cellular messengers [20, 21] involved in liver cell metabolism, contraction of ventricular, atrial, vascular, lymphatic and smooth muscle fibers, fluid secretion in intestinal, pancreatic, salivary, and sweat cells, aggregation of blood platelets, ion channel opening in T cells and astrocytes, differentiation in osteoblasts, proliferation of smooth muscle, brown fat, mesangial and T cells, fertilization, and neuronal synaptic plasticity in Purkinje and hippocampal neurons [22–27]. Calcium ions are stored in the endoplasmic reticulum (ER) at high concentrations (> 1 mM) while the concentration of free calcium ions in the surrounding medium (the cytosol) is tightly maintained at lower concentrations (100-700 *µ*M) [28]. Calcium is released from the ER by second messengers to raise intracellular Ca^2+^ concentrations and trigger a cellular response. Inositol trisphosphate, InsP_3_, is a ubiquitous second messenger involved the calcium-mediated processes listed above. It triggers Ca^2+^ release by binding and activating ion channels in the ER membrane [22, 29]. Sensitivity to InsP_3_ is modulated by the concentration of free intracellular calcium through calcium-induced calcium-release (CICR). CICR amplifies the signal so that a few InsP_3_ binding events can trigger a rapid release of calcium from the ER into the cytosol. Free calcium then diffuses through the cytosol. If enough calcium is released, then the diffusing calcium binds to an activating channel receptor and triggers the release of more calcium producing propagating waves [29]. Calcium also binds to an inhibitory channel receptor which inhibits further channel openings when the free calcium concentration is high [30]. Then the free calcium is gradually reabsorbed into the ER.

The combination of high InsP_3_ sensitivity at low Ca^+2^ concentrations, inhibition at high concentrations, and diffusion, allows complex temporal and spatial dynamics. These dynamics range from stochastic “blips” released by individual InsP_3_ channels, to “puffs” released by a cluster of channels, to travelling waves [28]. Shuai et al. present a discrete-state continuous-time Markov model for a single subunit of the channel to which InsP_3_ binds (hereafter, “InsP_3_ channel”). The subunit has nine states and 26 directed edges [30] (Figure 1). The subunits can change configuration, but are conserved in number, so Shuai’s model is an example of a reaction network model subject to a conservation constraint.

**Fig 1.**
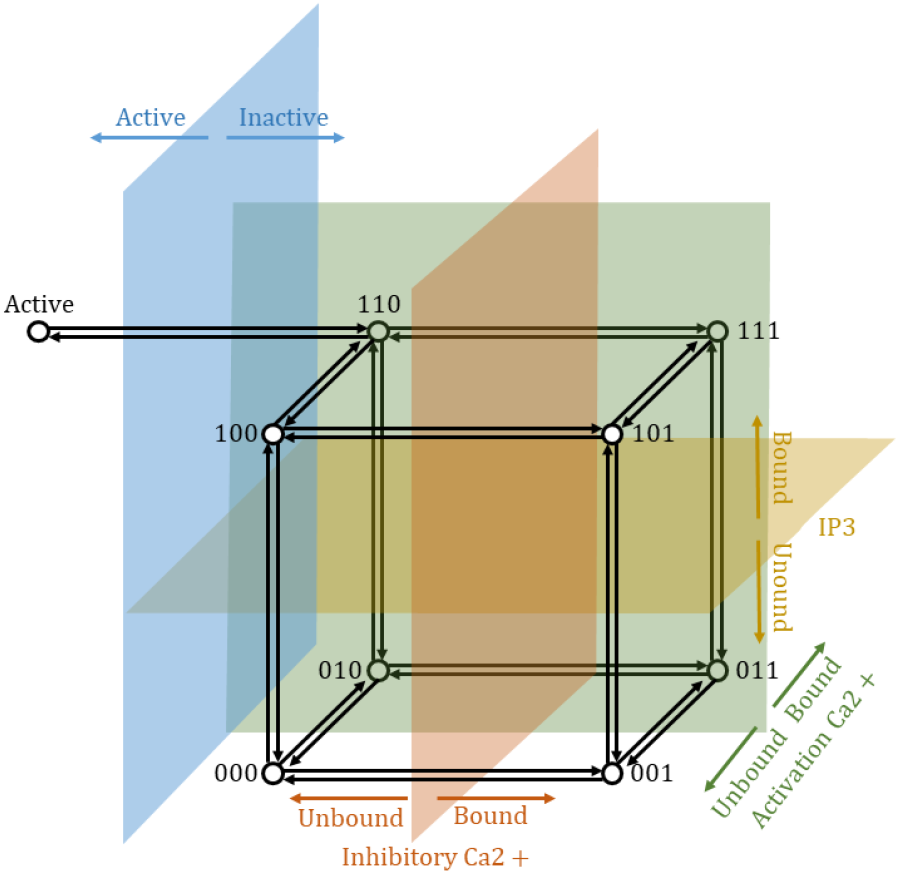
A single subunit in the InsP_3_ channel. The subunit has 9 states (vertices), 1 of which is active, and 8 which are inactive. The 8 inactive states form a cube where each axis corresponds to binding or unbinding at one of three different sites. Moving up binds inositol; moving front to back binds activation calcium, and moving left to right binds inhibitory calcium. In order to transition into the active state the subunit must bind to InsP_3_ (move up), bind activation calcium (move back), and not bind inhibitory calcium (move left).

One of the nine subunit states comprises an “active” configuration. The full InsP_3_ channel model contains four identical subunits, and the channel opens if at least three of the subunits are in the active state. As shown in Figure 1, each subunit has three binding sites, one for InsP_3_, and two for Ca^+2^. One calcium binding site activates the channel, and the other inhibits the channel. The first is responsible for CICR, while the second is responsible for closing the channel when free Ca^+2^ concentrations are high. Following [30], we use a 1 to denote a bound site and a 0 to denote an unbound site. Each subunit has nine states, eight corresponding to the 2^3^ possible states of the binding sites, and an additional state representing a conformational change that activates the subunit. The first eight states form a cube, and binding events move along the edges of the cube (black arrows in 1). The final state is connected to the (110) corner, since the subunit can only activate if it is bound to InsP_3_, the active calcium binding site is bound, and the inhibitory site is unbound. Shuai et al. demonstrated that, for appropriately chosen parameters, this model correctly reproduced the probability that the channel is active, and the expected time the channel spends closed as a function of calcium concentration [30]. The exact model and parameters are provided in [30]. Note that all binding events are first order reactions, and occur with rates that depend on the InsP_3_ and Ca^+2^ concentrations.

Since the number of active subunits controls whether a channel opens or closes, the only observable transitions are transitions into and out of the ninth state (active state). Since the model conserves the total number of subunits, it admits a steady state distribution and falls into the fourth limiting scenario (“persisting”). We study the importance of each noise source to the steady state variance in the number of open channels as a function of the InsP_3_ and Ca^+2^ concentrations. All importance values are calculated using Eq. 29. Finite time importance can also be computed using Eq. 11. Finite time importance could be relevant when considering the channel’s response to a spike in InsP_3_ or Ca^+2^. We focus on steady state importance here.

To compute importance, we specify a set of noise sources. We start by considering individual edges as noise sources; however the edges could also be grouped by reaction type. The importance of a reaction type can be recovered by summing the importance of the edges in that class. Four pairs of binding and unbinding edges correspond to the InsP_3_ binding site. Similarly, four pairs of edges correspond to binding and unbinding of the activator calcium binding site, and four pairs correspond to binding and unbinding at the inhibitory calcium site. These sets correspond to all edges of the cube that move up and down (InsP_3_), forward and back (activator calcium), and left and right (inhibitory calcium). The remaining pair of reactions corresponds to the conformational change from inactive to active (edges linking (110) to the active state). Since this pair contains the only observable reactions, the stochastic shielding heuristic would predict that these edges are the most important sources of noise for the number of active subunits [2]. However, we will show that the InsP_3_ and Ca^+2^ concentrations determine which edges are most important, and the pair of activating and inactivating reactions are not the most important in all situations.

The importance can be simplified by grouping forward and backward edges. The network contains 26 edges, consisting of 13 forward and backward pairs. The importance of the edges in each pair are always identical since the model obeys detailed balance [30]. In detailed balance, the steady state flux on all forward reactions matches the steady state flux on their backward partners. Moreover, if a forward reaction has stoichiometry *s*_→_ = *s* then the corresponding backwards reaction has stoichiometry *s*_←_ = −*s*, so 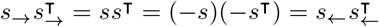. Therefore, the variance contributed by each forward reaction to the steady state will match the variance contributed by its backwards partner.

Importance is not evenly distributed among the 13 reaction pairs. For any combination of InsP_3_ and Ca^+2^ concentrations, the 4 most important edges account for more than half the steady state variance, and the 8 most important account for more than 90% of the steady state variance. The number of reactions needed to account for 50%, 90%, and 95% of the steady state variance is shown in Figure 2. Three of the four edge pairs involving the active calcium binding site never contribute more than a percent of the total variance. Binding of activating Ca^+2^ only matters if the inhibitory site is unbound, and the InsP_3_ site is bound. Most of the edges contribute less than a percent of the total variance when the calcium concentration is low (< 0.1 *µ*M) or the InsP_3_ concentration is low. Consequently, most edge pairs do not contribute significantly to the steady state variance unless the calcium concentration is intermediate (1 to 10 *µ*M) and the InsP_3_ concentration is small (< 1 *µ*M). Similar results extend to the classes of edges corresponding to particular reactions. Figure 3A shows regions in the Ca^+2^ InsP_3_ plane where each class of reactions is the least important.

**Fig 2.**
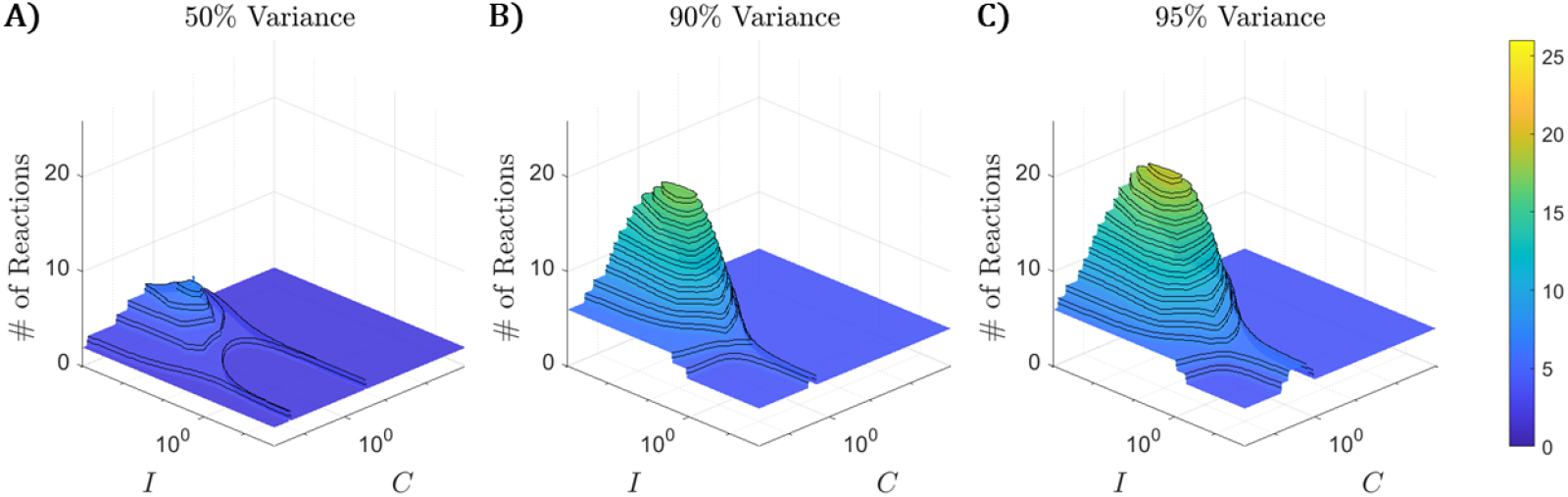
The number of edges needed to reach 50%, 90%, and 95% of the total variation in the number of active subunits depends on the InsP_3_ concentration (marked *I*) and the free Ca^+2^ concentration (marked *C*). All concentrations are in *µ* M and range from 10^−2^ to 10^1.75^ *µ* M. We never need more than 19 of the 26 edges to reach 95% of the total variance, and do not need most of the edges unless the Ca^+2^ concentration is intermediate and the InsP_3_ concentration is low.

**Fig 3.**
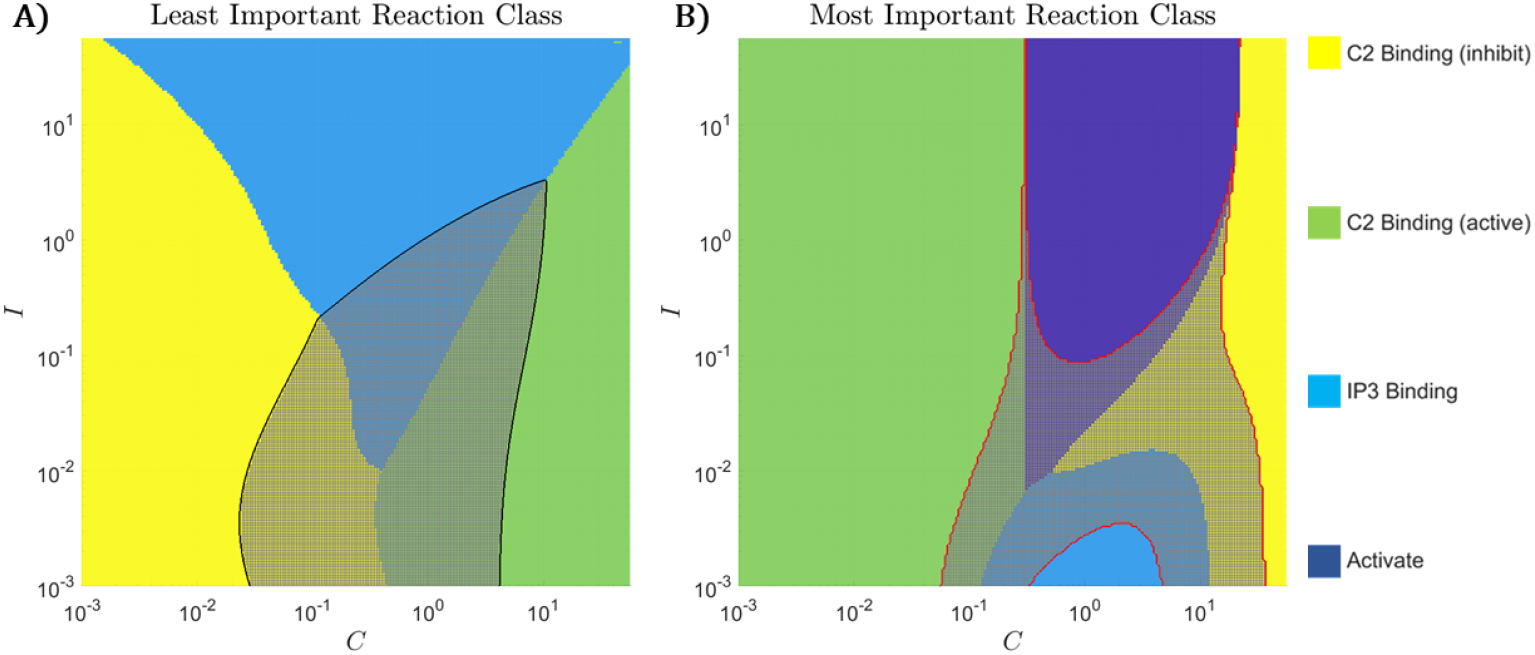
Importance maps. **A)** Reaction type contributing the least variance. The color of the region denotes which reaction type is least important. Grey hatching marks a region where the least important class contributes more than a percent of the total variance. **B)** Reaction types contributing the most variance. The color of the region indicates which reaction is most important. Grey hatching marks a region where the single most important class contributes more than half the total variance. The red line marks the boundary of this region, and matches the red contours shown in Figure 4.

Since most of the reactions do not contribute significantly to the variance in the number of active subunits the simulation of the InsP_3_ channel could be streamlined by simulating unimportant reactions deterministically rather than stochastically. If the free Ca^+2^ and InsP_3_ concentrations are held fixed for the duration of the simulation, then the importance of each reaction could be computed a priori, and unimportant reactions could be identified and simulated deterministically. This approach is easily implemented for Langevin approximations (c.f. [1, 7]) to the reaction network, and reduces the computation cost by reducing the total number of random numbers needed to perform the simulation.

Figure 3B shows regions where each class contributes a plurality and majority of the steady state variance. Binding to the activation site is most important when the Ca^+2^ concentration is low (< 0.1 *µ*M), and binding to the inhibitory site is most important when the Ca^+2^ concentration is high (> 10 *µ*M). Within the intermediate Ca^+2^ region, the conformational change into the active state is most important when the InsP_3_ concentration is not small (> 0.1 *µ*M), and the InsP_3_ binding site is most important when the InsP_3_ concentration is small (< 0.01 *µ*M). Thus, the directly observable reactions are only most important in the concentration regime where the channel spends most of its time activated. Outside of this region, the most important edges do not directly change the observable. This situation is an example of edge importance reversal [7].

Edge importance reversal occurs when the subunit spends most of its time closed, and only activates after a rare fluctuation. For example, the InsP_3_ binding edges are most important when the Ca^+2^ concentration is intermediate, but the InsP_3_ concentration is low. Then the subunit spends most of its time in the (010) state, and is immediately ready to activate if it happens to bind to InsP_3_ (move from 010 to 110). These binding events are rare when the InsP_3_ concentration is small. In contrast, if the subunit happens to bind to InsP_3_ then it moves to the 110 state, and is likely to activate (the activating and inactivating rates connecting the 110 to the active state are relatively fast). Then the rare InsP_3_ binding events contribute the majority of the variance in the number of active subunits.

Figure 4 shows the importance of the four classes of reactions. Each of the three classes of unobservable reactions are of negligible importance for a region corresponding to the colored regions in Figure 3A. In contrast, the importance of the observable reactions does not approach zero anywhere, and is never less than 0.14 (high Ca^+2^, low InsP_3_ concentrations). The observable reactions are never of negligible importance since the observable only changes when observable reactions occur. Thus, even if other reactions are more important, the observable events still matter.

**Fig 4.**
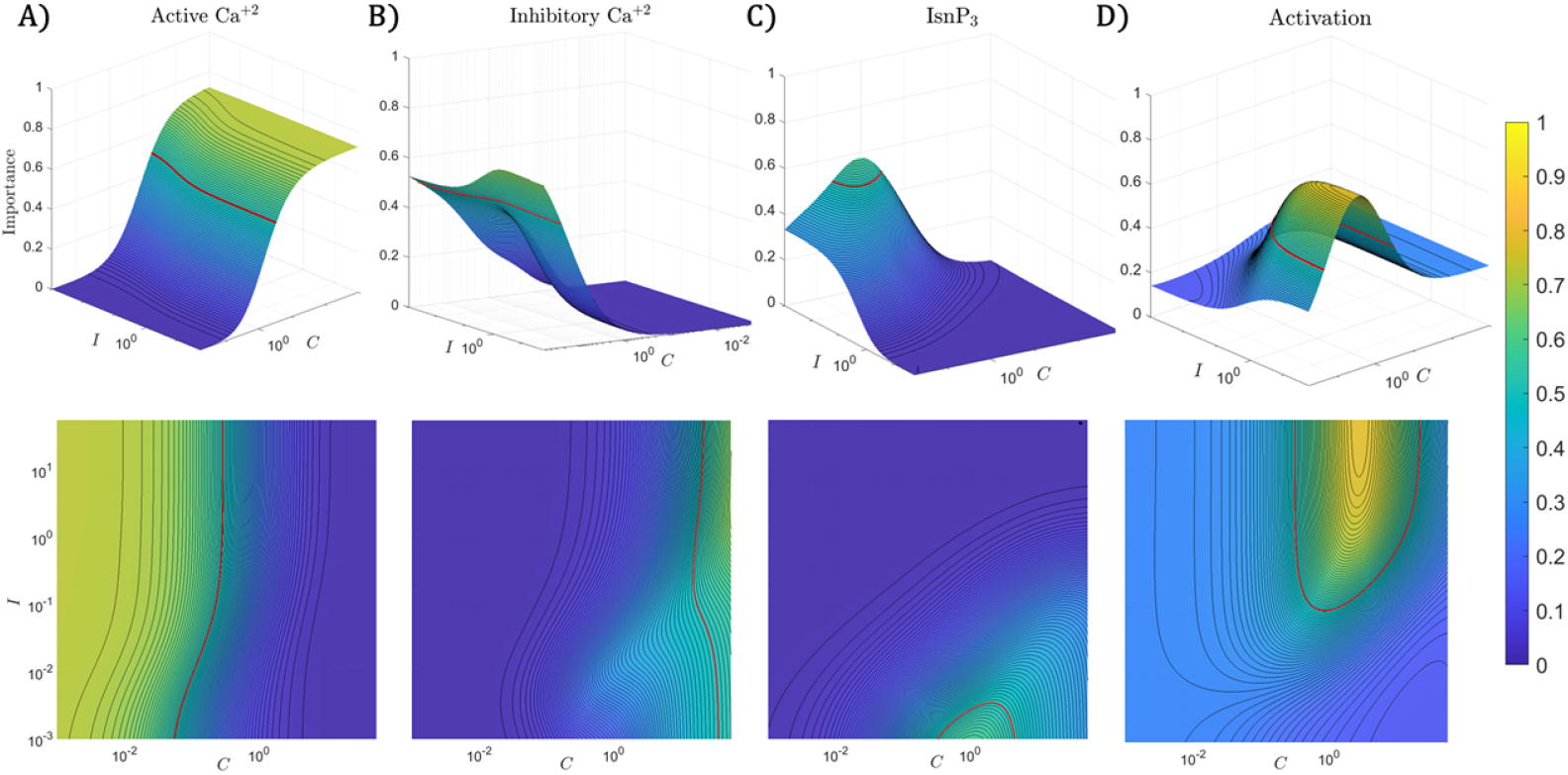
Importance surfaces. **A)** Importance of binding and unbinding to the activation site. **B)** Importance of binding and unbinding to the inhibition site. **C)** Importance of InsP_3_ binding and unbinding. **D)** Importance of the observable reactions. The bottom row shows each surface as a heat map. All panels share the same color bar. The red lines mark importance 0.5. Note that the importance of activation is largest when the Ca^+2^ concentration is small, and decreases as the concentration increases. Binding to the active site goes beneath 0.5 at the boundary where the green region ends in Figure 3B. Binding to the active site becomes unimportant when the Ca^+2^ concentration is large, near the boundary of the yellow region in Figure 3B. The importance of the inhibitory reaction is large when the Ca^+2^ concentration is large, and decreases as it decreases. Like the activation reaction, the inhibition reaction’s importance falls beneath 0.5 at the boundary of the yellow region, and becomes negligible in the green region of Figure 3B. This leaves an intermediate Ca^+2^ concentration regime (between 0.1 and 10 *µ*M) where the other two reaction classes are important. The InsP_3_ binding reaction is important when the InsP_3_ concentration is low, while the observable reactions (conformational change in the subunit between active and inactive) is important when the InsP_3_ concentration is large. The importance of the observable reactions closely follows the probability that the channel is active.

### Discrete-Time Matrix Models for Populations

Next, we present two examples of discrete-time matrix population models. The models incorporate age and spatial structure. Models of this kind are widely used in ecology to manage populations and to inform conservation efforts. *Population viability analysis*, among other methods, commonly uses discrete-time matrix models to guide management choices, determine which life events are most important for the success of the population, and the expected behavior population [14, 31]. Here we show that the tools developed in §Theoretical Results can be used to analyze noise structure in discrete-time matrix models. We identify which life events (e.g. reproduction, survival, dispersal, sex determination) produce the most variability in the population size at long times. We also identify when (at what age) and where (which transition) noise is most important. The examples are chosen to illustrate the theory. The first example offers growing and shrinking populations and has linear moment dynamics. The second example is a linearized nonlinear age and spatially structured metapopulation model. The linearization produces affine moment update equations, so requires the fully general analysis presented in §Affine Conditional Moments. We consider importance at both long times and at short times during a recolonization process.

### White-Tailed Eagles (*Haliaeetus albicilla*)

White-tailed eagles are a large, long-lived species found across the Palearctic. Like other species of eagle, white-tailed eagles suffered severe population declines in the 20th century due to direct persecution and pollution from heavy metals and pesticides such as DDT [32]. Eagle populations have recovered rapidly since the 1980’s. The eagles have been carefully studied in Germany since their recolonization in 1947. Both population and individual data are available for time series covering 62 years [33].

Here we consider a pair of discrete-time matrix models for white-tailed eagles in Schleswig-Holstein based on a model formulated by Krüger et al. A full description and parameterization can be found in [33]. Here we develop a stochastic interpretation of Krüger’s model that incorporates demographic stochasticity according to natural choices of distributions. Where needed, we estimate conditional variances using data in [33].

Krüger’s model is an age-structured model containing 37 age classes (ages 0 through 36 years). All individuals of age 36 are assumed to die. Eagles begin breeding after age four. Separate life history rates were estimated for the periods 1947 to 1974, when the species was endangered due to pollution, and 1974 to 2008, when the species recovered.

We assume that eagle deaths are independent, and that within a given time period (1947-1974 or 1974-2008), the probability an eagle dies depends only on its age. Then the number of eagles who survive in a given age class each year is a binomial random variable, with parameters fixed by the survival rate for that age class, and the number of individuals in that age class. The expected number of surviving birds from each age class is linear in the number of birds in the preceding age class, as is the variance. This model is supported by [34] who found no evidence for density dependence in eagle survival rates.

Survival rates were estimated by [33] based on age of death for 80 birds between 1953 and 2008. Due to small population sizes only 25 dead birds were recorded before 1974. As a result, survival rates were only estimated up to age 4 for the 1947-1974 period, and all survival rates for older eagles were set equal to the estimated survival rates for 1975-2008 [33]. The impacts of DDT on egg, chick, and adult survival rates are studied in [32, 35, 36]. Not all survival rates could be estimated directly. Survival rates for age classes missing data were set to the average of survival rates for neighboring age classes.

Here we present results based on Krüger’s reported survival parameters [33]. These parameters appear noisy, likely due to limited sample size. In order to evaluate the impact of parametric noise on our results, we also considered a version of the model in which survival rates as a function of age are smoothed after age 5. Survival rates were only smoothed after age 5, for two reasons. First, chick mortality is higher than juvenile mortality. Second, mortality at age 4 is high due to territorial competition between immigrant eagles of age 4 seeking a nesting site and adult eagles with established nests who defend their territories [34, 37]. We found that the smoothed model generated smoother trends in importance across ages, but as our qualitative conclusions were unaffected, we present results for the original unsmoothed model here.

We modeled the birth process as follows. The probability a particular female of a given age breeds is provided by [33]. Breeding probability is lower for young adults (0.620 at age 5) and approaches 1 for older adults (0.970 by age 8). Assuming independence, the number of females of a given age who breed is a binomial random variable with parameters equal to the probability any female breeds, and the number of females of the given age. Krüger et al. also supply the average fecundity of each age class, that is, the average number of fledged chicks per breeding female. Before 1975 each breeding female produced, on average, only 0.2 chicks each year. In contrast, after 1975 breeding females produced 0.55 to 0.77 chicks per year, with fecundity reaching a peak at middle age, before declining in older eagles. This dramatic change in fecundity is the main change in eagle life-history rates responsible for the recovery of the species [33].

To study the importance of each noise source we need an estimate for the variance in fecundity (fledged chicks per breeding female). Unfortunately, this variance is not provided explicitly in [33]. We used a two-pronged approach to bridge this gap. On one hand, we used clutch size data from [33] to estimate the variance. We validated our estimates against studies of Lithuanian and Swedish white-tailed eagle populations (see [38, 39]). On the other hand, we developed a variational approach that provided interval estimates for the importances. In the variational approach, we impose biologically motivated constraints on an unknown distribution, fix its mean, then maximize and minimize its variance given the constraints. We use this approach to avoid introducing specific distributional assumptions that are convenient but lack good biological motivation. For example, we avoid assuming that clutch sizes are Poisson distributed, as is assumed in some models (c.f. [34]), since clutch sizes above three or four eggs are virtually never observed and are physiologically implausible. Instead, we assumed that eagles: *a)* lay at most four eggs, *b)* lay zero eggs with probability 0.7 for 1947-1974 and 0.2 for 1975-2008 [33], and *c)* lay 3 eggs more frequently than they lay 4 eggs, as was observed in empirical clutch size distributions [38, 39]. Then, for each age class, we found the distribution with largest and smallest possible variance satisfying constraints *a*-*c* with mean equal to the average fecundity provided in [33]. The variances estimated from clutch size data [33] always fell between the max and min variance provided by the variational approach. Therefore, we report results from the variational approach which demonstrate the range of plausible importances across the range of plausible fecundity variance.

Eagle sex ratios at birth are close to 0.5 [33], so we assume that the number of female fledged chicks is a binomial random variable with mean equal to half the number of fledged chicks. We do not model males.

Each step in the life cycle is a separate noise source. We can then decompose the variance produced by birth into components associated with separate life events. Since we only track the number of fledged female chicks produced at the end of the birth sequence, we use the law of total variance to decompose the variance in the number of fledged female chicks into contributions from each random event (whether a female breeds, clutch size, and sex determination). The sum of these contributions is the overall contribution of noise from reproduction. Decompositions of this kind could be introduced to other steps where there is enough modelling information to decompose the events. For example, the number of fledged chicks could be separated into the number of eggs laid, then the number of eggs that survive. Mortality could also be decomposed into mortality from separate causes (disease, predation, starvation, etc.) given enough data. The actual structure of the decomposition into separate noise sources is both model and species dependent. For example, some species of birds double-clutch (c.f. [40]), in which case the birth event should include a separate random event that accounts for the number of clutches laid in a breeding season.

Pre and post 1974 models differ primarily in fecundity. Fecundity before 1974 was reduced by egg-shell thinning due to pollution, and resulted in a declining population. Our model, like Krüger’s, predicts a 6% annual decline in eagle population each year prior to 1974 (*λ*_1_ = 0.947). After 1974 the model predicts a 4% annual growth in the eagle population (*λ*_1_ = 1.04). Both the growth and decay rates are consistent with Krüger’s predictions [33].

To compute noise source importance, we set the initial population equal to ten individuals, and partition the population into age classes according to the stable age distribution. We then compute the importance of each noise source to the total population using equations 23 and 24. Figure 5 illustrates results for the maximum variance case.

**Fig 5.**
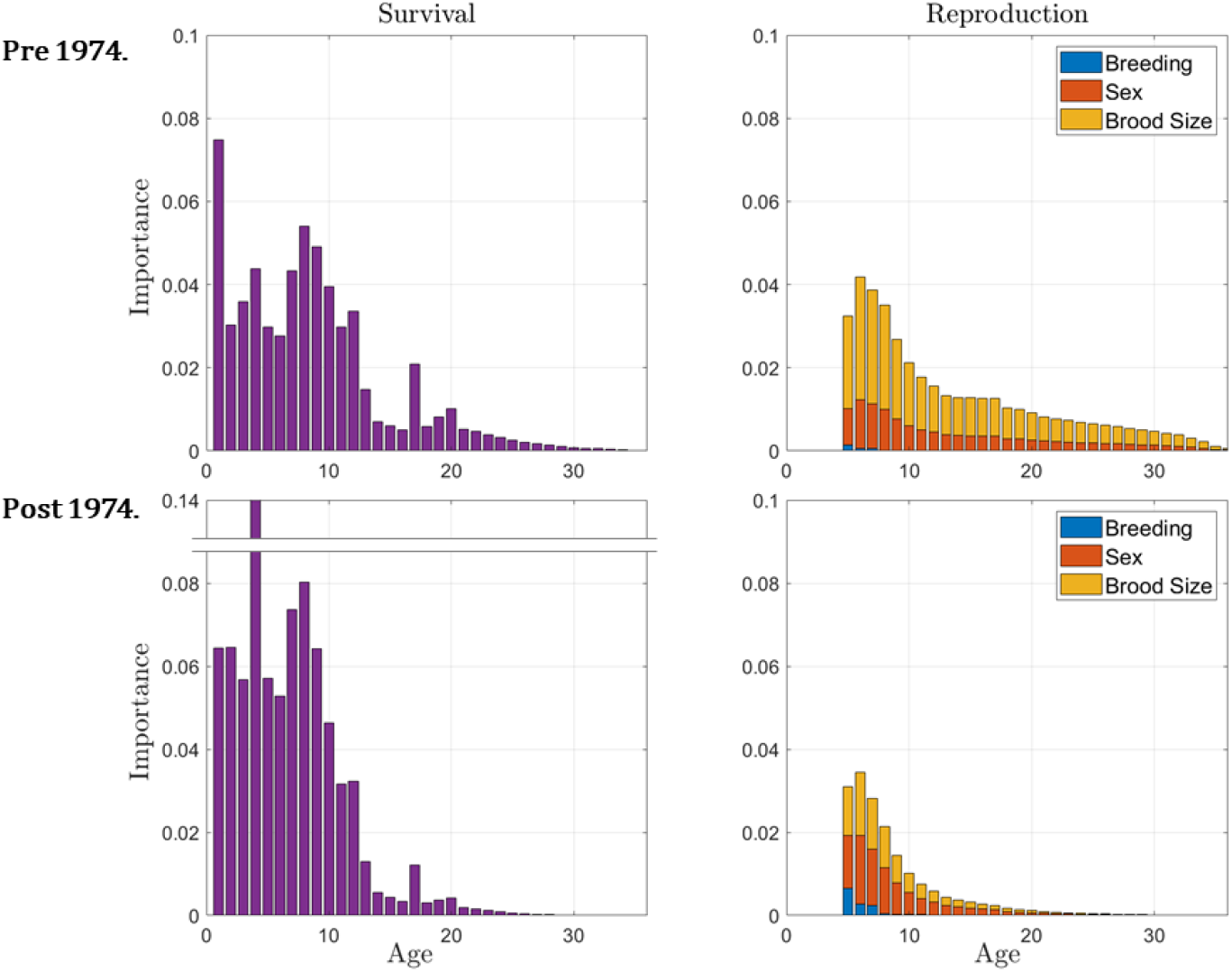
**Top row:** Importance of noise sources for variance in white-tailed eagle populations before 1974, assuming maximum variance in clutch sizes. Note the high variance from chick mortality, large noise contributions from ages 4 and 8, and the slow decay in the importance of reproduction. The erratic behavior of importance in mortality at later ages is likely due to noise in the estimated survival rates, and was smoothed out after smoothing the survival rates (not shown). **Bottom row:** Importance of noise sources for white-tailed eagles after 1974, assuming maximum variance in clutch sizes. Note the spike in importance at age 4, preceding the transition to sexual maturity, and for young adults experiencing greater mortality.

In all cases, we find that mortality contributes more to the overall noise in eagle populations than reproduction, with mortality accounting for 59 to 62% of variance before 1974 (max and min variance in clutch sizes), and 79 to 82% of all variance after 1974. After 1974 the importance of mortality peaked at age 4. This peak can be explained in two ways. First, eagle mortality is high at age 4 due to territorial contests between young adults hoping to establish a new nest, and established adults. Notably, this increase in mortality is not observed before 1974 when eagle populations were small, so nesting space was less limited [33]. Second, the transition from age 4 to age 5 is crucial, as it represents the transition to sexual maturity. Before 1974 the most important noise source is chick mortality. We find that, across most population models studied, the most important noise source is either mortality preceding the transition into sexual maturity, or mortality during the first year of life, which is often high. In both eras, we see large noise source importance for mortality during the first 8 to 10 years of life. These are crucial years, since individuals must survive the first four years to reproduce, have notably lower survival rates as young adults, and take approximately 8 years before they breed reliably.

Reproduction accounts for 38 to 40% of the total variance in population size before 1974 (min and max variance in clutch sizes), and 18 to 20% of the total variance after 1974. The variance contributed by whether or not a female breeds is largest at age 5 and declines quickly as 97% of all females older than 8 breed. Thus, uncertainty in breeding generally contributes the least variance of all noise sources considered (≤ 1%), and is negligible for eagles over 8 years old. Before 1974, variance due to clutch size is consistently larger than variance due to individuals’ sex, accounting for 26 to 29% of the total variance. Variance due to individuals’ sex accounted for 11 to 12% of the variance in these cases. After 1974, clutch size and sex contribute comparable amounts of variance, accounting for 8 to 12% of the total variance each. In general, the importance of each reproduction event peaks at age 6, and then declines steadily with increasing age.

Accounting for all sources of noise at a given age we find that, before 1974, noise from the first year of life is particularly important, and total importance peaks at age 7. That said, noise at later ages is still important. The first 10 years of life account for 62.5% of the total variance, the first 15 years account for 78.9% of the total variance in population size, and the first 20 years account for 89.4% of the total variance. Similar results hold after 1975, except noise source importance peaks at age 4, and declines faster, with the first 10 years of life accounting for 83.6% of the total variance in the eagle population.

Thus, when eagle populations are shrinking, reproduction, in particular clutch size, contributes more to the total variation in the population, importance is distributed more evenly across age classes, and adults in their prime reproductive years (near age 8) and chicks contribute most of the noise. When eagle populations are growing, reproduction contributes notably less of the total variance, importance peaks at age 4 (the transition into sexual maturity), and decays faster with age.

Note that, to perform this type of analysis we do not need a full stochastic population model, only the conditional moment update equations. Full stochastic models are helpful when the distribution of a random quantity can be inferred naturally from the life-history. For example, as long as deaths occur independently, survival can usually be modeled as a binomial random variable, in which case the conditional mean and variance are parameterized by the survival probabilities. However, we are often missing information needed to build a full stochastic model. For example, birth events are often more difficult to model, since clutch, brood, or litter sizes do not necessarily fit a standard family of distributions. Instead annual reproductive success often depends on a sequence of random events, resulting in distributions that are often multimodal or skewed [16]. Nevertheless, it is possible to use empirical information to constrain the moment update equations as illustrated by our variational approach. In all cases, had we assumed a Poisson or binomial distribution for clutch size we would have overestimated the variance in clutch size.

### Wood Frogs (*Rana sylvatica*)

Wood frogs are a well-studied (c.f. [41–43]) North American amphibian. In order to study the relative importance of distinct noise sources within wood frog populations, we linearize a nonlinear metapopulation model. We consider two distinct ecological scenarios: equilibrium across life stages and local habitats, and global recovery from a large perturbation. Because the linearized conditional moment equations are affine, we apply the results from §Affine Conditional Moments. In addition, this example demonstrates how to analyze noise source importance along transients, in particular during a colonization process following a large perturbation.

Adult wood frogs migrate to ponds to breed in the early spring, where adult females lay a single egg mass. There, the eggs hatch into fully aquatic larvae which metamorphose into terrestrial juvenile frogs in the early summer. Wood frogs typically reach sexual maturity after two years on land before they return to the pond to breed as what we will refer to as “young adults”. Some frogs, which we call “mature adults”, survive to breed a second time in the following year. When wood frog ponds are spread across a large terrestrial habitat, they may act as a metapopulation [42, 44]. While the majority of individuals return to breed in the pond they hatched in, some frogs disperse during their juvenile life stage and breed in a new pond [42]. Variation in dispersal contributes to population variability within individual ponds, and as a whole. Thus, unlike the eagle model, the wood-frog model incorporates spatial structure and noise from dispersal.

The source wood frog metapopulation model (Rollins, Benard, Huffmyer, and Abbott, in prep.) was a nonlinear deterministic ODE model based on data collected by Michael F. Benard at the University of Michigan’s E. S. George Reserve (ESGR) shown in 6. The model focuses on females, operates on a discrete one year time scale, includes 21 ponds and 4 age classes: eggs, juveniles, young adults, and mature adults. The continuous terrestrial environment surrounding the ponds is shared by all juvenile and adult individuals. The model is nonlinear since, as in other amphibians, survival from egg to juvenile is density-dependent (due to larval competition within each pond), as is survival from juvenile to young adult (reflecting juvenile competition across the whole terrestrial environment) [43, 45, 46]. Each of these density-dependent transitions uses a Beverton-Holt model [47], which outperformed linear versions of each transition in AIC [48] goodness of fit tests. Note that the model ignores certain important features of the ESGR by assuming that the ponds are equivalent, thereby ignoring the differences in predation between large and small ponds. For these reasons, our results should not be interpreted as specific predictions about the ESGR, which would require a more detailed model, but as illustration of our theory in a model with realistic age, stage, and spatial structure.

To make the model amenable to our measures, we replace it with a stochastic model whose conditional moment equations are affine. We perform this conversion in two steps. First, we define a fully stochastic model consistent with the deterministic nonlinear model, and compute the conditional moment equations. These equations are nonlinear due to density dependence in survival. Next, we fit a linear stochastic model to long simulated trajectories from the nonlinear stochastic model. Advantages and disadvantages of this approach, as well as other linearization methods, are discussed in S4 Appendix

The stochastic model follows. First, all young and adult female frogs reproduce in each pond. While the form of the egg mass distribution is not known (roughly log-normal), the necessary moments can be estimated from available data. Each female lays 723.7± 197.3 eggs (mean and standard deviaiton). For the sake of simulation, we use a log-normal distribution, rounded to integer values. We assume that the number of eggs laid by each female is independent, so the variance in the total number of eggs laid in a pond is the sum of the variance in the number of eggs laid by each female. We do not consider sex at birth, or the probability a female does not reproduce, because data on tadpole sex is not available and females who did not return to the pond habitat to breed cannot and need not be distinguished from females who had died. Egg survival to the juvenile stage is binomial with a density dependent survival probability:

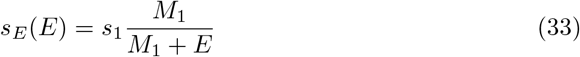

where *s*_1_ is the survival probability of a single egg in the absence of any other eggs, *M*_1_ is the half-saturation constant, and *E* is the number of eggs in the pond. Surviving eggs become aquatic tadpoles, who metamorphose into terrestrial juveniles.

Juveniles then enter the terrestrial phase, during which time they disperse. Juveniles are terrestrial for two years, so we introduce a lag population that counts the terrestrial juveniles who have not yet reached sexual maturity.

Next we model dispersal. Each juvenile individual has a 94% chance of returning to their birth pond to breed once they mature, so only a small fraction disperse. The remaining frogs disperse according to an exponentially decaying dispersal kernel. The probability of a juvenile dispersing from ponds separated by distance *d* is set proportional to exp(− *d/d*_0_) where *d*_0_ is a reference distance (average dispersal distance of frogs that do not return to their natal pond). Note that this dispersal model does not distribute frogs equally across the ponds. Instead, well connected ponds tend to receive more immigrants than they produce emigrants, so have higher total steady state populations. Since survival is nonlinear and the ponds don’t maintain the same total populations, the ponds also don’t share exactly the same age-distribution, though dispersal is rare enough that the effect on the age distribution is small.

Once a breeding pond has been assigned to each juvenile, we sample a fraction that survive to arrive in that pond at the end of the two-year juvenile stage. Survival probability *s*_*L*_ is density dependent, and depends on the total number of individuals in the terrestrial environment. Specifically,

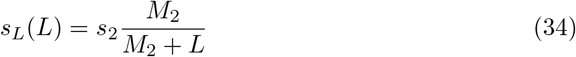

where *L* is the total lag population (individuals in the terrestrial environment that have not yet reached breeding age), *s*_1_ is the survival probability for an individual in absence of any others, and *M*_2_ is a half-saturation constant. The number of young adult individuals that survive to breed in pond *i* is then sampled from a binomial distribution with survival probability set to *s*_*L*_(*L*) and number of trials set to the number of lag individuals assigned to migrate into pond *i*.

Young adults who survive another year return to their breeding pond one year later as mature adults. Survival of the young adults is not density dependent, so the number of mature adults each year is drawn from a binomial distribution with a fixed survival probability *s*_*Y*_. All mature adults die in the subsequent year.

To validate the stochastic model, we ran the deterministic ODE model to equilibrium, and compared the equilibrium to the average populations in each age class and pond after a long (2× 10^5^ years) trial run of the stochastic model. We found that the deterministic equilibrium agrees with the average predicted by the stochastic model, to within a maximum relative error 0.6% over all age classes and ponds.

The stochastic model does not have linear conditional moments, because egg and lag survival are both density dependent (see equations 33 and 34). So, to apply the noise source importance measures, we require a linearization procedure. We linearize by repeating long realizations (2 ×10^5^ years) of the nonlinear stochastic model, estimating the conditional moments, and fitting to affine functions.

After linearizing about the steady state, the conditional moment updates for the number of juveniles *J*_*t*_(*i*) in pond *i* at time *t*, given the number of eggs *e* in the pond, are 𝔼[*J*_*t*_(*i*) |*E*_*t*_(*i*) = *e*] = 0.0136*e* + 938.3 and Cov[*J*_*t*_(*i*) |*E*_*t*_(*i*) = *e*] = 0.0133*e* + 905.3. Ninety-five percent confidence intervals for the four parameters are [0.0134, 0.0138],[926, 951],[0.0122, 0.0145], and [840, 970]. Note that the slope was usually determined with much higher precision than the intercept, since both linear fits are primarily sampled at large egg counts.

The number of young adults who arrive at pond *k* depends on both the survival rate of frogs during the pre-reproductive (“lag”) terrestrial phase, which depends on the total lag population, and the number of surviving lag individuals who choose pond *k*. Thus, we fit the expectation and variance in the number of immigrating lag individuals into each pond to the number assigned to that pond and the total number of lag individuals. The associated fits were 𝔼[*Y*_*t*+1_(*j*)|*L*_*t*_ = *l*] = 0.031*l*(*k*) − 0.00065 Σ_*i*_ *l*(*i*) + 22.9 and Cov[*Y*_*t*+1_(*k*)|*L*_*t*_ = *l*] = 0.027*l*(*k*) − 0.0008 Σ_*i*_ *l*(*i*) + 34.37, with confidence intervals, [0.307, 0.031], [− 0.0007, −0.0006], [20.7, 25.1], [0.024, 0.030], [−0.0015, −0.0002], and [11.3, 57.4].

Once the conditional mean and variance of each intermediate stage have been approximated, we compute the contribution of each noise source by using the law of total variance to group sequences of reactions that account for the transitions between tracked variables.

With the linearized model established, we compute the importance of each noise contribution (egg counts, egg survival, dispersal, juvenile survival, and young survival) to each age and space class (the total frog population, the number of juveniles in each pond, number of young in each pond, number of adults in each pond, and number of reproductive individuals (young and old) in each pond). We also tested the importance of each noise source to the total frog population in each pond. All noise sources were identified by a life history event, producing 105 different noise sources. Figure 7 illustrates the results.

First, we observe that the importances depend much more strongly on which life event is considered, than which pond it occurs in. Figure 7 illustrates this effect: the variance in importance between life history events (differences among box heights) is much larger than the variance across ponds (size of box and whisker associated with each event).

Broadly speaking, the most important event is juvenile survival, followed by young adult survival, egg survival, egg count, then dispersal, though the importance order may change depending on which life stages are observed. For example, egg survival and count are more important to variance in juvenile population size than young adult survival is, but young adult survival is more important than any other event, even juvenile survival, to variance in the mature adult population. Juvenile survival is the most important event in most cases since it is the survival stage preceding reproduction. Dispersal is the least important in all cases since, in the linearized model, the ponds are indistinguishable and interact only with the total lag population, which is independent of pond assignment. Consequently, while dispersal contributes to the variance in the number of frogs in any individual pond, it does not contribute any variance to the total population across ponds. Dispersal also plays a small role since the 94% of all frogs return to their birth pond.

So, dispersal (and the associated spatial component of the model) is not an important source of randomness in the process if we consider the total population across all ponds after reaching a stationary distribution. However, if we consider specific ponds, or a transient away from equilibrium, then dispersal plays a more important role.

For example, if only a single pond is considered, then all five noise sources associated with that pond are more important than noise generated in other ponds, and match the ordering of importance shown for the total population across ponds. The importance of noise sources in the other ponds is generally very small. If we track the population in pond *i*, then the most important event in pond *j* is always dispersal, since the only way noise can propagate from pond *j* to pond *i* is via dispersal. This fact is an elegant illustration, in an ecological context, of stochastic shielding [1, 2, 7].

Dispersal is also important during transients. During particularly dry summers, some of the ponds in the reserve may desiccate, killing off the aquatic life stages. In subsequent, wetter years, frogs will disperse out from the larger ponds to repopulate smaller ponds that dried out. To complete our study, we considered the importance of each different noise source over the course of an recolonization from the largest pond in the reserve. The largest pond in the reserve is Fish Hook Marsh (Pond 8 in Figure 6). We initialize the system by setting the frog population to 0 in all ponds except pond 8, and set the population in pond 8 to its multi-pond equilibrium value. This initial condition is far from equilibrium, so we need to relinearize the model. S4 Appendix describes the relinearization technique.

**Fig 6.**
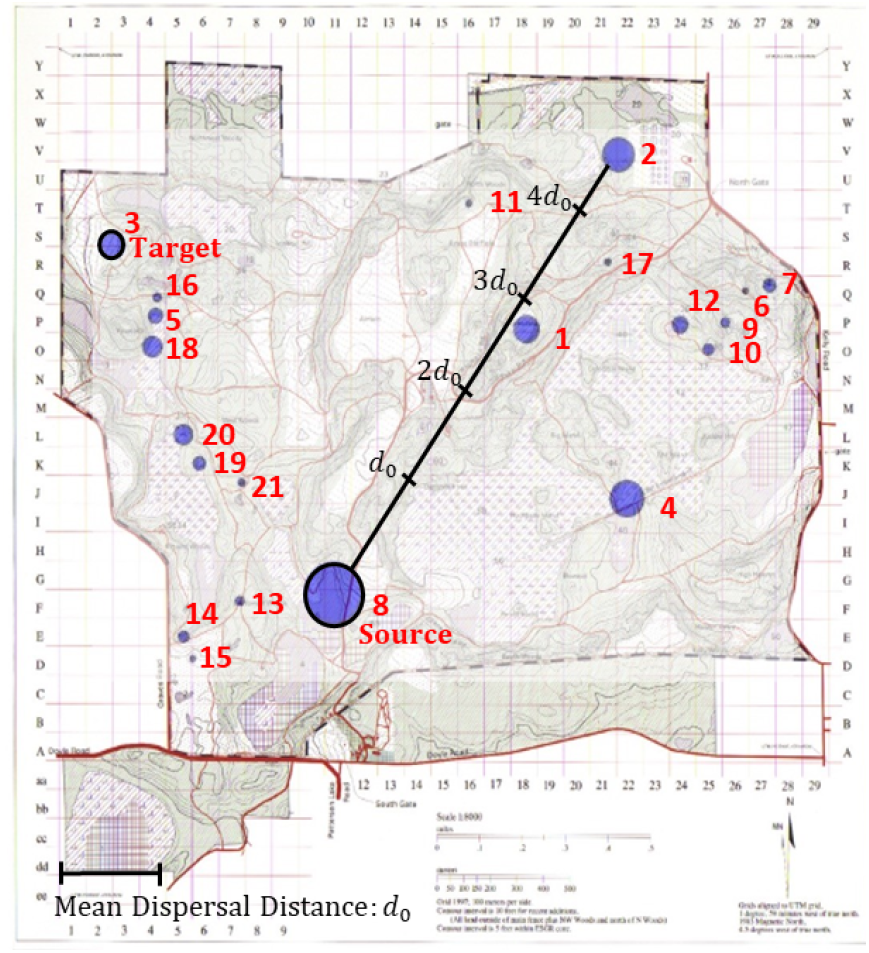
Map of the 21 ponds in the E. S. George Reserve, adapted from [49]. The ponds are numbered, and relative sizes of the ponds are represented by the sizes of the blue circles marking each pond. Pond 8 (Fish Hook Marsh) is significantly larger than the other ponds, and ponds 1 - 4 are at least 3 times larger than the remaining ponds. The mean dispersal distance *d*_0_ is marked with a solid black line on the bottom left. The distance from pond 8 to pond 2 is shown in units of *d*_0_ for scale. All ponds within a distance of 4.6 *d*_0_ are expected to exchange one disperser each year when the source pond population is at its local equilibrium. We report equilibrium results in which all ponds are occupied, as well as results for colonization from pond 8 to pond 3 (marked source and target respectively).

**Fig 7.**
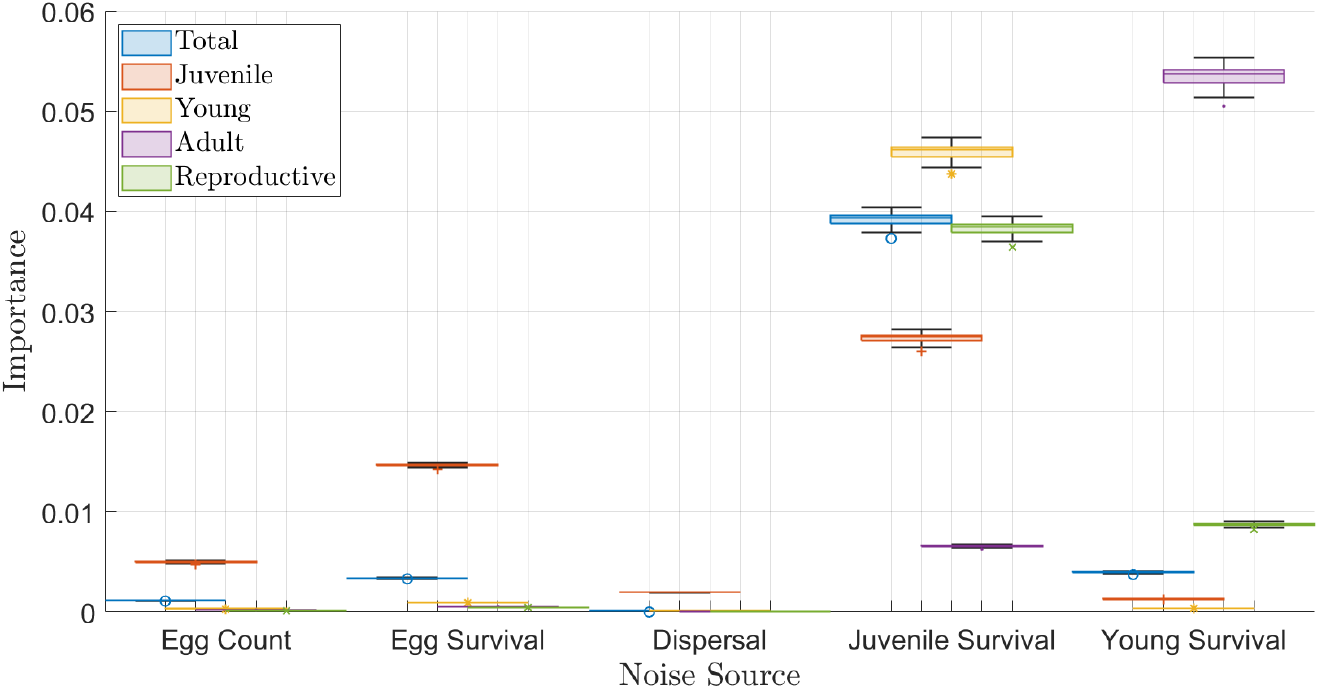
Importance of each noise source for five different observables. The five observed quantities are the total frog population (blue), juvenile population predestined for each pond (orange), young adult population in each pond (yellow), mature adult population in each pond (purple), and the total population of reproductive individuals (young + mature adults) in each pond (green). The noise sources are organized by life-history event. The box plot for each life history event represents the distribution of importance across the 21 ponds. The horizontal central line is the median importance, the box marks the first and third quantiles, whiskers mark the range excluding outliers which are marked with individual points.

After linearizing about the perturbed initial conditions, we computed the importance of each noise source to the total population in a set of target ponds. Figure 8 shows the results when the source pond is set to pond 8 and the target pond is set to pond 3. This example is representative. Pond 3 was chosen as the displayed target since it is connected to Pond 8 by a chain of smaller ponds, so, for the right dispersal parameters we might expect to see a cascade of importance in dispersal down the chain from 8 to 3. Pond 3 is also an interesting target since it is large, but far removed from 8, and neighbors a cluster of smaller ponds (16, 5, 18).

**Fig 8.**
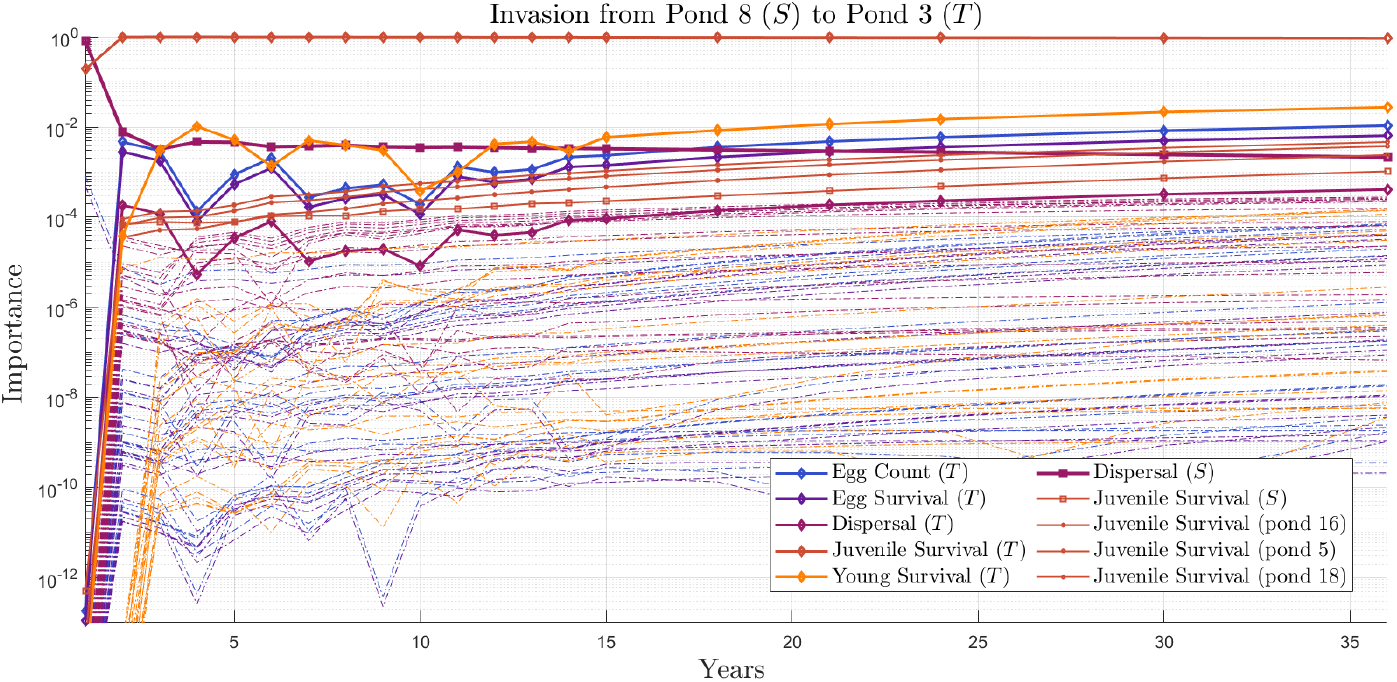
Importance of each noise source during recolonization from pond 8 (source *S*) to pond 3 (target *T*). The horizontal axis represents the cumulative importance of each noise source up to a specific year since recolonization. Frogs live at most 3 years. Consequently, the model exhibits decaying oscillations with period 3 years. So, we sample the process at one year intervals from 1 to 15, and 3 year intervals from 15 to 36. Each line represents the importance of all the noise produced by a particular source up to each year. The lines are colored by life event. Light blue: egg count. Dark purple: egg survival. Magenta: dispersal. Rust orange: juvenile survival. Yellow: young adult survival. The noise sources shown with dashed lines are all negligible (importance less than 10^−4^), so could be ignored in a reduced stochastic model of the invasion process. The noise sources shown with solid lines are marked according to their pond, and labeled in the legend. Diamonds represent events in the target pond. Squares mark events in the source pond. Circles mark events in the closest neighbors of the target pond (ponds 16, 5, 18, in decreasing proximity to the target).

In the first year, the most important noise source to variance in the target pond’s population size (pond 3) is dispersal out of the source pond (pond 8). Dispersal out of the source pond remains among the most important noise sources in later years, but decays quickly in importance after the first year. Juvenile survival in the target pond replaces dispersal out of the source pond as the most important source in year two, then remains the dominant source of noise in all but the first year. As the frog population in the target pond grows, the importances of the other life events in the target pond grow, approaching the importance order seen at equilibrium. While the noise sources in the target pond are the most important sources at relatively long times, dispersal out of the source pond remains more important than most of the noise sources in the target pond (excluding juvenile survival) until 15 - 18 years after invasion. This slow decay in the importance of dispersal is noteworthy given how little dispersal, contributes at equilibrium.

The slow decay of the importance of dispersal out of the source after the first year is a result of the cumulative nature of the measure. Dispersal importance out of the source pond includes all variance contributed by dispersal out of the source pond up to the current year. Thus, since dispersal out of the source pond is necessary to seed the frog population in all other ponds, dispersal out of the source in year one is a crucial noise source that contributes a significant portion of the noise in the target pond at future times.

Juvenile survival in the source pond is also important for similar reasons. Dispersal is a relatively rare event, so the target pond may not be seeded during the first year post invasion. In that case, survival of the juveniles in the source pond is important for seeding the target pond in subsequent years.

The remaining important noise sources are all associated with the cluster of ponds neighboring the target pond (16, 5, 18). Juvenile survival in ponds 16, 5, and 18 are the next most important noise sources, and are ordered in importance by their distance from the target pond. Thus, spatial structure influences importance during the initial phase of colonization. This effect stands in contrast to the situation at equilibrium, in which spatial structure plays no role.

On the other hand, at intermediate time scales, importance rapidly shifts from dispersal out of the source to local dynamics within the target pond. Indeed, after the first time step, over 95% of the total variance in the target population is generated by variance in juvenile survival in the target pond. This rapid shift results from the relative time-scales of dispersal and the within-pond dynamics. While only a small fraction of individuals disperse (7/117 in the model realization depicted), the number of juveniles who survive to disperse is itself large enough (1749) that most of the ponds are seeded in the first time step. If the distance *d*_0_ which controls the average dispersal distance were small, then we would expect to see invasion spread in a travelling front across the reserve. We do not observe this effect here. The mean dispersal distance *d*_0_ is large enough that most ponds are seeded in the first year, consistent with the treatment of the terrestrial environment as a single well-mixed patch. In fact, all of the ponds are close enough to pond 8 so that the expected number of immigrating individuals after one time step is greater than 1. The initial seeding leads to rapid growth in the local population, at which point the within-pond dynamics produce most of the relevant noise, since the time scale for mixing through dispersal is slower than the time scale of the within-pond dynamics.

## Discussion

The measures developed here apply to a wide range of stochastic models used in biology. They reveal how noise propagates through those models, and they identify important and unimportant noise sources for a range of quantities of interest. In most of the examples explored here, a small subset of noise sources contributed most of the uncertainty in the quantity of interest, suggesting that noise source importance could be used to guide model reduction.

Not all stochastic models have linear, or approximately linear, conditional moment dynamics. Future work could investigate whether it is possible to generalize the importance measures developed here to other functional forms for the conditional moments. In particular, when applied to population models, our framework is largely limited to demographic noise, since environmental noise typically arises via time varying model parameters, which introduce multiplicative noise. In separate work [50], we have developed analogous noise source sensitivity analysis that applies to a class of discrete-time matrix models with time varying rates (c.f. [51]).

While we focused on case studies in this paper, it would be interesting to perform a meta-analysis of collections of stochastic models that share a linear formulation. For example, the COMADRE and COMPADRE databases [52, 53] could be mined for discrete-time matrix populations models as in [50]. A study of this kind could identify whether there are general patterns that predict noise source importance. Future theoretical work could also consider families of random networks, and search for network signatures which identify which noise sources are likely to contribute the most to a given observable.

## Supporting information

**S1 Appendix. Continuous-time analysis**. This appendix derives the continuous-time variance decomposition and noise source importance measures presented in §Theoretical Results.

**S2 Appendix. Convergence of discrete-time approximations**. This appendix shows that the noise source importance measure associated with a discrete-time approximation to a continuous time process converges to the noise source importance measure for the original continuous time process.

**S3 Appendix. Affine conditional moment equations**. This appendix derives the noise source importance measures presented in §Affine Conditional Moments.

**S4 Appendix. Linearization of the wood frog model**. This appendix documents the linearization procedures used to approximate the nonlinear wood frog model with a linear model. The methods are compared, and their respective advantages and disadvantages are discussed.

## Acknowledgments

This work was supported by NSF grants DEB-1654989 to KCA and PJT, DMS-1840221 to KCA, and DMS-2052109 to PJT. PJT acknowledges research support from the Oberlin College Library. We gratefully acknowledge Professor Michael Benard (CWRU) for providing access to data from his wood-frog population study.

